# Multiscale excitation-inhibition balance dynamics: integrating metabolite kinetics with time-varying executive networks

**DOI:** 10.1101/2024.10.30.621153

**Authors:** Francesca Saviola, Stefano Tambalo, Laura Beghini, Asia Ferrari, Barbara Cassone, Dimitri Van De Ville, Jorge Jovicich

**Affiliations:** CIMeC, Center for Mind/Brain Sciences, University of Trento, Rovereto (TN), Italy; Department of Medical and Surgical Specialties, Radiological Sciences and Public Health, University of Brescia, Brescia, Italy; Neuro-X Institute, Ecole Polytechnique Fédérale de Lausanne (EPFL), Geneva, Switzerland; Department of Physics, Faculty of Natural Sciences, Norwegian University of Science and Technology, Trondheim, Norway; Department of Clinical and Experimental Sciences, Neurology Unit, University of Brescia, Brescia, Italy; Department of Psychology, University of Milano-Bicocca, Milan, Italy; Department of Radiology and Medical Informatics, University of Geneva (UNIGE), Geneva, Switzerland

## Abstract

The balance between neural excitation and inhibition (EIB) is an essential mechanism supporting cognitive processes. Yet, little is understood about how EIB shifts with cognitive load and its impact on functional connectivity dynamics. In this study, we investigate temporal profiles of the reciprocal modulation between EIB and functional network dynamics during working memory tasks, revealing that EIB prefrontal kinetics scale with increasing cognitive load. Notably, prefrontal EIB kinetics correlated with cognitive load, impacting stability of networks crucial for cognitive function. On one hand, brain dynamics adapt to meet increasing cognitive challenges with a shift towards more focused and sustained neural activity patterns in terms of connectivity. On the other, imbalances favouring excitation may hinder cognitive adaptability. Importantly, this experimental approach demonstrates a link between EIB kinetics, brain network dynamics and cognitive performance, defining the groundwork for exploring healthy and aberrant cognitive states.

**Teaser:** Highly focused or less responsive? Chemical signalling and network dynamics are coupled to produce persistent cognitive states.

## Introduction

The functionality of the human brain hinges on a delicate balance between two principal neurotransmitters: glutamate, the primary excitatory neurotransmitter, and γ-Aminobutyric acid (GABA), with an inhibitory effect on neuronal excitability^1^. Their ratio, known as Excitation-Inhibition Balance (EIB), is crucial for maintaining cognitive functions and synaptic plasticity^2–4^. Disruptions in EIB are implicated in numerous neurological and psychiatric disorders, highlighting its critical role in both brain health and disease^5^. While substantial research has focused on the static roles of excitatory and inhibitory circuits, there is growing recognition of their dynamic interplay and its significance in human cognition^6^. To date, little is known about how EIB dynamics affect the functional connectivity of networks supporting cognitive processes.

Preclinical studies have demonstrated both the feasibility and the cognitive relevance of in-vivo EIB mapping^7–9^. However, non-invasive EIB estimation in humans remains challenging, pushing the field to advance through the development of computational approximation models. Indeed, these proxies probed how microscale human brain organization strongly hinges upon hierarchical computationally-derived EIB gradients^10–12^ with a strong association to blood oxygen level-dependent (BOLD) signal temporal variability^13^, both in health and disease^5,14,15^. All of the above stated the significance of the relationship between time-varying patterns of functional connections and EIB kinetics, which urge to be deepened through more reliable estimations of the latter.

Recent modelling approaches have revealed complex temporal dynamics of neurometabolites as measured by functional Magnetic Resonance Spectroscopy (fMRS). Two contrasting hypotheses emerged, each proposing distinct timescales for metabolite fluctuations. Lea-Carnall et al. (2023)’s model^16^ suggested rapid shifts in glutamate and GABA levels between vesicular and cytosolic compartments, occurring within seconds of neural activity changes. In contrast, Mangia et al., (2009)’s hypothesis^17^ argued for slower processes linked to energy metabolism and operating over several minutes. These divergent perspectives highlighted the multifaceted nature of fMRS signals, which likely encompass a spectrum of temporal dynamics from rapid neurotransmitter cycling to gradual metabolic adjustments. The relative contributions of these fast and slow processes may vary depending on brain regions and experimental conditions, underscoring fMRS data complexity.

This dynamic complexity is particularly relevant to understanding brain functions like working memory (WM), a core component of higher-order cognition and goal-directed behaviour. WM significantly relies on prefrontal cortices, where glutamatergic pyramidal cell firing persists to maintain WM function^18^ and GABAergic circuit disruptions can result in severe deficits^19^. There is currently no evidence of human dynamic regulation of EIB during WM tasks, and it remains unclear how changes in neurometabolic substrates relate to time-varying functional connectivity in prefrontal cortices.

To address this gap, we introduced an innovative experimental approach designed to investigate a key cognitive neuroscience question — WM modulation under varying cognitive loads — through the integration of functional neuroimaging (fMRI) and molecular mechanisms (fMRS). The novelty of the protocol is two-fold. First, it involved interleaving functional MRI with functional MRS^20–22^, enabling non-invasive examination brain connectivity and neurometabolites coupling as a function of cognitive load. Secondly, we adapted fMRS to estimate EIB kinetics for increasing cognitive load. We speculate that this protocol opens the path to applications in various domains, such as basic cognitive or clinical neuroscience research as well as validation and refinement of computational models that provide mechanistic links between neurobiological activity and behaviour in health and disease.

To demonstrate the feasibility of our protocol, we applied it to healthy young volunteers using a clinical 3T MRI system. Specifically, we explored the hypothesis^23^ that different mental workloads during WM tasks elicit parametric alterations of EIB kinetics, reflecting fluctuations in executive functional networks over time. Our first aim is to detect alterations in neurometabolites, especially excitatory and inhibitory ones, across sessions characterized by different levels of cognitive load. The second aim involves moving beyond static estimation of neurometabolites concentrations to investigate their cycling during the fMRS time course, offering insights into the timescale of responses. Lastly, our third objective is to compare and integrate the time-resolved contributions of metabolic and vascular information in sustaining goal-directed behaviour. To accomplish this, we employed time-varying models of functional connectivity and assessed associations with metabolic data.

## Results

### In-vivo detection of EIB kinetics

Out of 25 subjects with fMRS acquisitions, only 12 met the stringent data quality criteria. Indices of spectral quality and the group-averaged edited GABA+ and Glx spectra from the left DLPFC (grey matter fraction: 0.52±0.04%) are reported in the Supplementary materials for the four WM loads considered throughout the functional paradigm: Rest, 0-Back, 1-Back and 2-Back. Notably, static fMRS data analysis revealed no significant WM load effects on normalized concentration measures averaged during each cognitive condition. This lack of effects held true for GABA+, Glx, and EIB ratio, as determined by Kruskal-Wallis tests.

However, dynamic analysis yielded more nuanced results (Figure 1A). A Kruskal-Wallis test on EIB Area under the Curve (AUC) revealed a significant effect of Load (χ2(3,44)=9.95, p-value=0.02, Figure 1B). Post-hoc comparisons demonstrated that the 2- Back condition exhibited a significantly higher EIB curve increase relative to the resting-state condition (Rest > 2-Back, p-value=0.04, Mean Ranks Difference=-14.9, Figure 1B). Similarly, a significant effect was observed for GABA (χ2 (3,44)=10.64, p-value=0.01). No significant effects were found for Glx across different frames.

**Figure 1:**
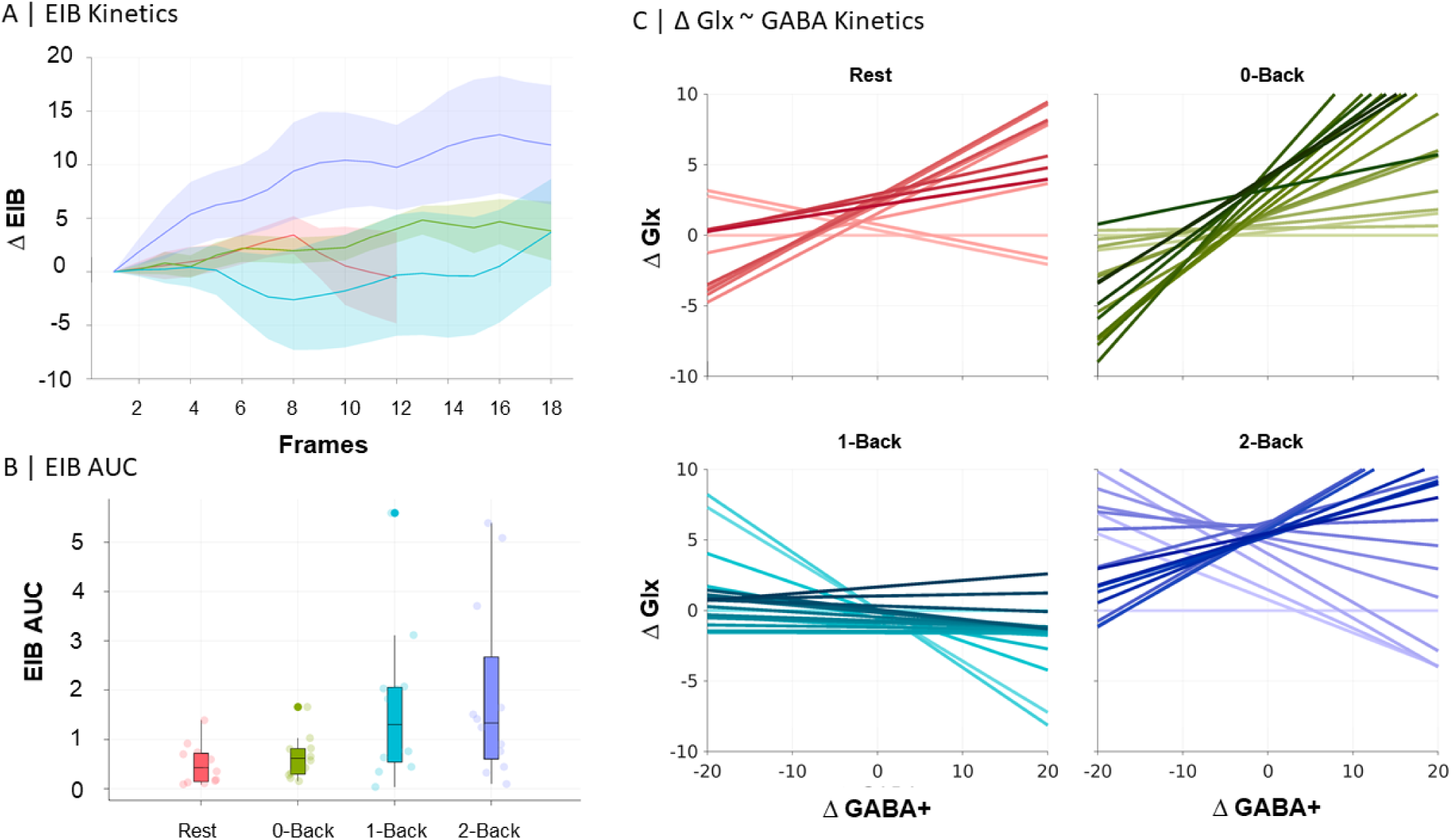
Changes of in-vivo EIB kinetics across cognitive load levels. A: Kinetics of EIB curves using a sliding-window approach (4 min per frame, with 12 seconds shifts) for each cognitive condition. Thick lines show the mean and the shaded areas the standard error of the mean. B: AUC Boxplot derived from A depicting Load effects between Rest and 2-Back sessions. C: Evolution of Glx/GABA+ imbalance to reveal the directionality of the effect. Temporal dimension is depicted as a color gradient from light (first frame) to solid (last frame).

These findings highlight the importance of monitoring the evolution of neurometabolite concentrations, as they reveal task-related modulations that are not apparent in static measures. The observed increase in EIB during the high cognitive load condition (2-Back) suggests a potential adaptive mechanism in brain metabolism to meet increased cognitive demands.

### Comparing executive network temporal dynamics and EIB kinetics

The FPN seed-based dynamic fMRI analysis identified four FPN-CAPs across all sessions, consistently in both the full sample of 36 subjects and a subset of 12 subjects with concurrent MRS acquisition. These FPN-CAPs, matched to the Yeo atlas24 using cross-correlation (r-value>0.3), exhibited distinct temporal patterns corresponding to varying cognitive loads (see Supplementary Materials). Table 3 summarizes the temporal properties of FPN-CAP networking for the full sample; Table S4 presents data for the MRS subset. Both reveal significant differences in higher cognitive load conditions compared to resting state (Figure 2). Notably, FPN-CAP temporal dynamics increased significantly as mental workload incrementally rose, stabilizing during higher cognitive task conditions.

**Figure 2:**
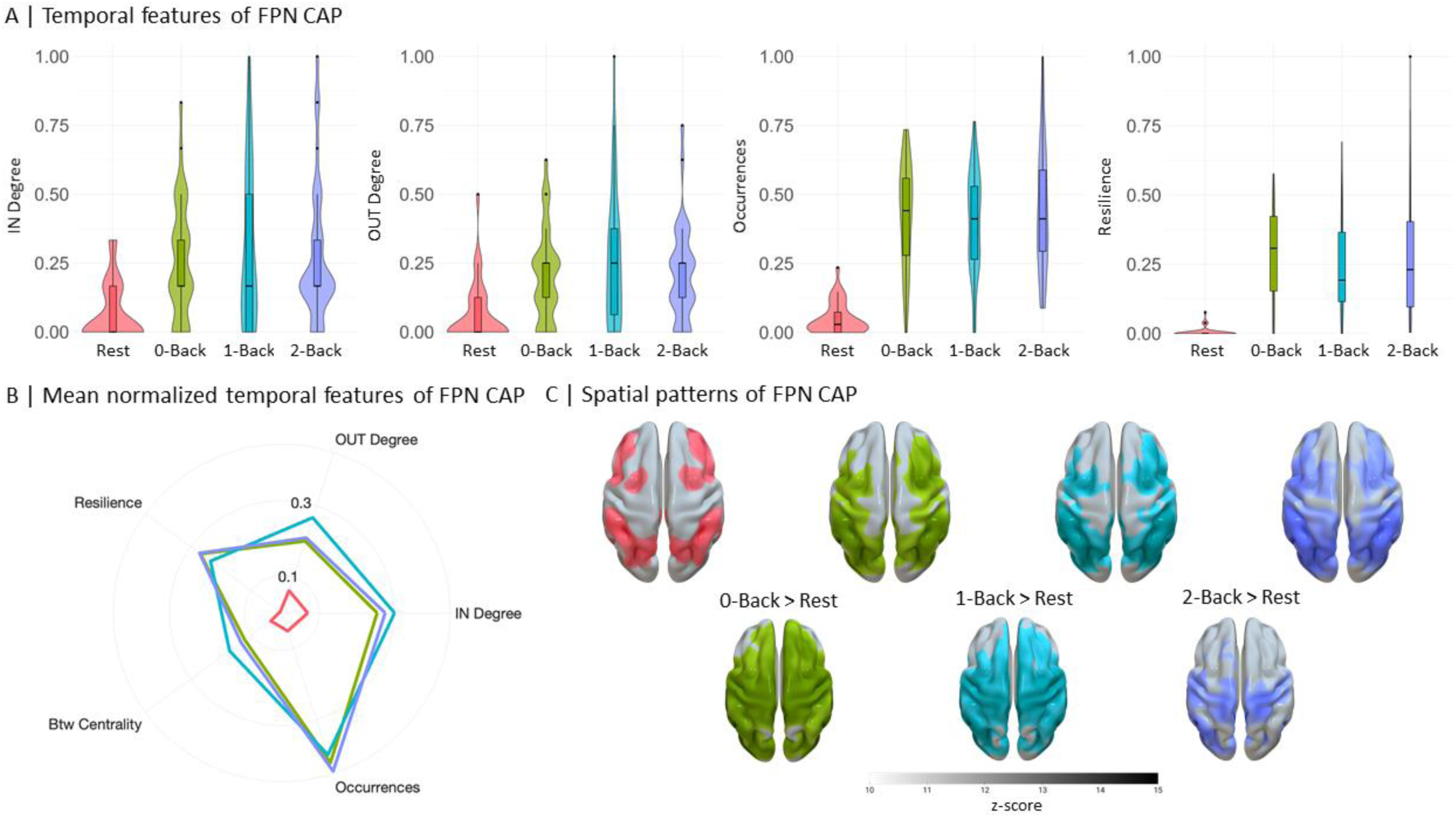
Executive networks temporal dynamics moving toward stationarity as a function of cognitive load. A: violin plots of normalized FPN-CAP temporal properties exhibiting significant changes across sessions (p<0.001) B: radar plot displays the level of temporal dynamic properties of FPN networking over the time-series. Temporal properties are represented as normalized scores ranging 0-1 with respect to data distribution across runs. C: Voxel-wise differences of back- projected FPN-CAP. First row depicts one-sample t-test results for the main effect of sessions. Second row shows pairwise t-tests across sessions (FWE across voxels and Bonferroni across comparisons p<0.016).

**Table 3:** Temporal properties of FPN different sessions in fMRI sample.

| Temporal FPN properties | Working memory conditions |  |  |  | Differences across conditions (FDR corrected) |  |  |
| --- | --- | --- | --- | --- | --- | --- | --- |
|  | rest | 0-back | 1-back | 2-back | Post-hoc comparisons | p-value | Chi-square/ A-B estimate |
| **Occurrences (%)** | 1.7±1.9 | 14.3±6.4 | 13.5±6.2 | 15.1±7.7 |  | **<0.001\*\*** | **69.4** |
|  |  |  |  |  | **0-back>rest** | **<0.001\*\*** | **66.9** |
|  |  |  |  |  | **1-back>rest** | **<0.001\*\*** | **62.5** |
|  |  |  |  |  | **2-back>rest** | **<0.001\*\*** | **69.1** |
| **Resilience** | 0.0007±0.002 | 0.04±0.02 | 0.03±0.03 | 0.04±0.03 |  | **<0.001\*\*** | **65.75** |
|  |  |  |  |  | **0-back>rest** | **<0.001\*\*** | **65.75** |
|  |  |  |  |  | **1-back>rest** | **<0.001\*\*** | **61.06** |
|  |  |  |  |  | **2-back>rest** | **<0.001\*\*** | **64.43** |
| **Betweenness centrality** | 0.2±0.4 | 0.6±0.9 | 0.9±1.3 | 0.7±1.2 |  | >0.05 |  |
| **IN Degree** | 0.002±0.003 | 0.008±0.006 | 0.009±0.009 | 0.008±0.008 |  | **<0.001\*\*** | **25.35** |
|  |  |  |  |  | **0-back>rest** | **<0.001\*\*** | **37.83** |
|  |  |  |  |  | **1-back>rest** | **<0.001\*\*** | **39.00** |
|  |  |  |  |  | **2-back>rest** | **<0.001\*\*** | **38.94** |
| **OUT Degree** | 0.003±0.004 | 0.008±0.007 | 0.010±0.010 | 0.009±0.007 |  | **<0.001\*\*** | **25.93** |
|  |  |  |  |  | **0-back>rest** | **<0.001\*\*** | **35.98** |
|  |  |  |  |  | **1-back>rest** | **<0.001\*\*** | **42.87** |
|  |  |  |  |  | **2-back>rest** | **<0.001\*\*** | **37.83** |

Spatial topographies of FPN-CAP also showed significant voxel-wise changes across sessions (p-value_FWE_<0.016, Bonferroni corrected). The activation pattern evolved from anatomically confined areas during resting state to a more extensive distribution under higher cognitive loads. In the 2-Back condition, this expanded pattern encompassed all key FPN hub regions, with notably increased involvement of lateral prefrontal areas. Pairwise comparisons further confirmed these findings, revealing larger differences between resting-state activations and those during attentive (0-Back) or lower load (1-Back) conditions. This progression underscores the FPN’s adaptive engagement in response to increasing cognitive demands.

To integrate neurovascular information from dynamic functional connectivity with kinetics of neurometabolic concentrations, we analysed visibility graph-based time-series properties of the EIB curve. Specifically, we focused on out-degree (expressing a temporal gradient of AUC) and Kullback-Leiber divergence (KLD; index of temporal reversibility) across all conditions (Table S5). A one-way ANOVA across sessions revealed statistically significant differences in the out-degree of EIB (χ2 (3,44)=9.92, p-value=0.02). Post-hoc comparisons showed an increased out-degree of EIB in the 1-Back condition compared to the resting state (Rest<1- Back, p-value=0.02, Mean Ranks Difference=-16.5). This suggests a change in the temporal dynamics of EIB as cognitive load increases.

While no significant differences were found in the KLD, we observed an increasing trend as a function of cognitive load. The comparison between Rest and 2-Back conditions returned a qualitative, statistically non-significant trend (Rest<2-Back, Mean Ranks Difference=-12.37, p=0.06) towards temporal irreversibility in EIB under higher cognitive demands.

These findings highlight the complex relationship between cognitive load and EIB kinetics: the increased out-degree in the 1-Back condition suggests more frequent changes in EIB levels, while the rising KLD trend in higher cognitive loads points to time-dependent changes in EIB. Together, these results suggest that neurometabolic processes and functional network reorganization are adaptive and reciprocal, responding dynamically to varying cognitive demands.

### Behavioural performance results

Behavioural measures (i.e., RTs, accuracy and *d’*) revealed a significant effect of load over sessions (p<0.05, see Supplementary Materials) for all variables of interest. This enabled the idea that, as previously shown in literature^25^, the level of cognitive load was increasing across sessions and therefore brain functional connections and neurometabolism should change accordingly.

### The effect of EIB kinetics and executive networking on behavioural performance

Our analysis revealed complex relationships between neurometabolites, FPN temporal properties, and task performance. The Generalized Linear Model (GLM), which analysed behavioural performance as the dependent variable and included various fixed-effect factors such as session, block, and participant demographics, highlighted a significant role of static GABA+ concentration in predicting RTs (β = -0.1, p-valueFDR < 0.001; see Supplementary Materials). However, neurometabolites kinetics showed no significant effect on behavioural performance. Although previous research has established a correlation between elevated GABA levels and improved task performance as measured by accuracy^26,27^, our findings do not demonstrate a significant relationship between accuracy and GABA+. Instead, the observed significant association with RTs implies that higher GABA+ concentrations may be linked to enhanced readiness and attentional capacity during task execution, which is manifested as faster RTs.

In contrast, FPN-CAP temporal properties demonstrated significant associations with cognitive performance. Three key dynamic features emerged as performance predictors, both individually and in interaction across sessions: in-degree (likelihood of entering an FPN state), out-degree (likelihood of exiting an FPN state), and occurrences (frequency of a given state). In particular, for *d’*, we observed significant effects for out-degree as a fixed effect (β = 7.9, p- value_FDR_ < 0.001) and in interaction with 1-Back (β = -18.6, p-value_FDR_ < 0.001) and 2-Back sessions (β = -5.2, p-value_FDR_< 0.001). Accuracy was significantly affected by out-degree interacting with 1-Back session (β = -13.9, p-value_FDR_< 0.001), in-degree interacting with both 1-Back (β = -9.8, p-value_FDR_< 0.05) and 2-Back sessions (β = -11.5, p-value_FDR_< 0.05), and occurrences interacting with both 1-Back (β = -0.005, p-value_FDR_< 0.05) and 2-Back sessions (β = -0.005, p-value_FDR_< 0.05) (see Supplementary Materials). Notably, no significant effects were found for RTs. The negative relationship between behavioral performance and FPN-CAP temporal properties suggest that during the 1-Back and 2-Back runs, task performance may decline as FPN-CAP properties increase. This instability within the FPN-CAP may imply that heightened engagement of the network could lead to variations in cognitive control and attentional resources, which might contribute to challenges in maintaining optimal task performance.

Looking more closely at the relationship between FPN-CAP temporal properties and static neurometabolite concentrations, we identified significant effects exclusively for GABA+. All assessed FPN-CAP temporal properties significantly predicted GABA+ levels, particularly in interaction with the 0-Back session. Specifically, we observed significant interactions for out-degree (β =-0.2, p-value_FDR_< 0.01), in-degree (β =-0.2, p-value_FDR_< 0.01), occurrences (β =-0.2, p-value_FDR_< 0.01), betweenness centrality (β =-0.2, p-value_FDR_< 0.01), and resilience (β =-0.2, p-value_FDR_< 0.01) (see Supplementary Materials).

### The cognitive modulation of temporal dynamics and its top-down effects on EIB kinetics

To deepen our understanding of the relationship between neurometabolic kinetics and FPN- CAP temporal features across WM cognitive loads, we conducted a Partial Least Square Correlation (PLSC) analysis. This analysis focused on the subset of subjects with MRS measurements (N=12), utilizing visibility graph properties of EIB curves. For this purpose, we performed the CAP analysis by concatenating data from all 12 subjects across different sessions, using the left DLPFC as a consistent seed region (see Supplementary Materials). PLSC was chosen for its ability to incorporate multimodal brain properties, finding optimal linear combinations of CAP temporal features and EIB kinetics features that maximally correlate with each other.

The PLSC analysis yielded a significant Latent Component (LC) as determined by permutation testing (covariance explained = 59%, p=0.03). This LC captured the greatest covariance between FPN temporality and EIB kinetics. Figure 3B illustrates the maximization of correlation between brain scores achieved by the analysis (r=0.43, p=0.002).

**Figure 3:**
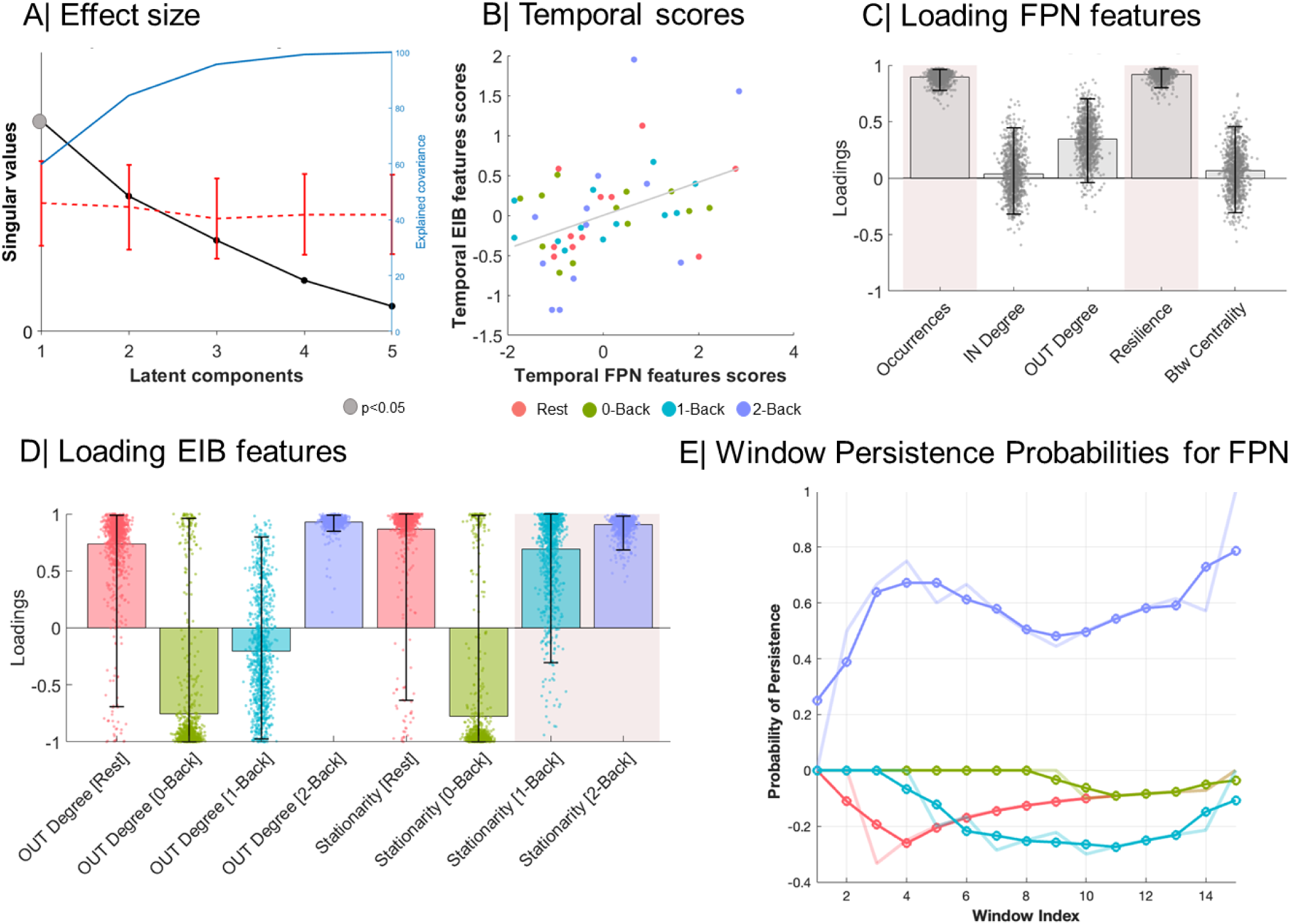
Dynamics of neurometabolites concentration contribute to constrain temporal reconfiguration of cognitive states. A) Partial least squares analysis (PLS) was used to assess the multivariate relationship between EIB kinetics temporal features and executive network (FPN) temporal features. PLS identified a single significant latent variable (Covariance explained=59%, dotted p-value=0.03; two-tailed). Line plot describes the median, bounds represent the first (25%) and third (75%) quartile of the distribution. B) Temporal patterns of EIB and FPN scores are depicted for the first Latent variable. The two brain score patterns are significantly correlated (p=0.0022, r=0.43; two-tailed). R-values denote the Pearson correlation coefficient, linear regression line collapsed across groups is added to the scatterplot for visualization purposes only. C) PLS loading FPN features are related to all temporal properties of executive networking and are depicted as bar charts with their corresponding 95% confidence interval from 10000 bootstrap resampling. Pink boxes describing reliable loading are plotted for significant bootstrap testing. Spatial distributions of executive CAP networking are depicted across the cortex for each cognitive workload condition. D) PLS loading EIB features are shown for each workload condition as bar charts (a single PLS loading per EIB temporal scores for each session) with their corresponding 95% confidence interval from 10000 bootstrap resampling. Pink boxes describing reliable loading are plotted for significant bootstrap testing. E) fMRS-matched sliding-window approach to investigate time scale similarities of FPN-CAP with EIB kinetics. Abbreviations: Fronto- parietal network (FPN), Co-activation pattern (CAP), Betweenness Centrality (Btw), Excitation-inhibition balance (EIB).

We then examined the corresponding FPN-CAP and EIB kinetics temporal feature loadings across sessions, identifying the features that contributed most significantly to the spatial patterns captured by the first LC (Figures 3C, 3D). This integrated analysis approach provides a comprehensive view of how FPN dynamics and EIB kinetics co-vary across different cognitive loads, offering insights into the neural mechanisms underlying WM processes. The significant LC and strong correlation between brain scores underline the tight coupling between network dynamics and neurometabolite fluctuations during cognitive tasks hypothesized in the previous section.

Our analysis revealed that temporal properties of EIB kinetics showed diverse correlations with FPN dynamics across sessions, with a particular emphasis on occurrences and resilience, suggesting their potential role in sustaining neurotransmission. The temporal gradient of the EIB curve (represented by average EIB Out-degree) and its stationarity (interpreted as the inverse of KLD) exhibited characteristic associations with FPN temporal properties. As cognitive load increased parametrically, both EIB Out-degree and EIB stationarity initially showed negative associations with temporal FPN properties, primarily driven by occurrences and resilience^28^. However, this relationship shifted towards a positive association under higher mental workload conditions.

To summarize, our analysis revealed that brain weights were particularly robust for EIB kinetics in terms of stationarity across sessions. This finding indicates that the corresponding pattern of correlation weights was heavily influenced by the KLD of EIB. As subjects progressed through different cognitive states, the EIB curve became less time-invariant (i.e., less stable) for higher workload conditions. As cognitive demands increase, the brain appears to transition from a more stable EIB state to a more flexible one, potentially allowing for greater adaptability in neural processing.

To gain deeper insights into the temporal scale similarities between the two measurements, we analysed the probability of persistence for the given FPN-CAP across EIB-matched time intervals (i.e., window indices; Figure 3E, see Supplementary Material for other CAPs). This analysis revealed intriguing patterns of CAP stability across different cognitive loads.

A Kruskal-Wallis test on persistence curve points demonstrated a significant effect of Load (Chi-squared(3,56)=38.9, p-value<0.001, Figure 3E). Post-hoc comparisons yielded particularly interesting results for the 2-Back condition, which showed a significantly higher window’s persistence probability curve increase compared to other conditions: (i) Rest>2-Back (p-value<0.001, Mean Ranks Difference=-32.2); 0-Back>2-Back (p-value<0.01, Mean Ranks Difference=-20.2); 1-Back>2-Back (p-value<0.001, Mean Ranks Difference=-35.2). These findings suggest that the 2-Back condition, representing the highest cognitive load in our study, is associated with a distinct pattern of FPN-CAP persistence. The increased persistence probability in this condition may indicate a more stable or sustained engagement of the relevant brain networks during high-demand cognitive tasks. This observation provides valuable insights into how brain dynamics adapt to meet increasing cognitive challenges, potentially reflecting a shift towards more focused and sustained neural activity patterns under higher cognitive loads.

## Discussion

At the current state-of-the-art, EIB is thought to play a major role in many leading theories for the pathophysiology of brain disorders, especially neurodevelopmental and psychiatric ones^5^. In this respect, the time-dependent mapping of EIB enabled by the usage of fMRS represents a unique window on the dynamics of neurochemical signalling in the human brain.

Seminal studies^24,25^ have shown the feasibility to map EIB with fMRS. Craven et al.(2024)^21^ revealed that glutamate and GABA levels fluctuate on different timescales during cognitive tasks, with glutamate exhibiting more rapid changes (seconds to minutes) compared to GABA (minutes to tens of minutes). Moreover, Mullins et al.(2024)^26^ showed that these temporal dynamics of neurometabolites can vary depending on brain region and task complexity, highlighting the intricate nature of EIB regulation. Expanding on these findings, Emeliyanova et al. (2024)^27^ introduced a novel perspective by providing evidence for the existence of distinct "*metabolic states*" that transition on timescales of seconds to minutes during cognitive processing. This discovery suggests a more complex temporal structure to EIB than previously conceived, bridging the gap between rapid neurometabolic fluctuations and broader cognitive states.

However, a clearer description of the temporal interplay of excitatory and inhibitory chemical drivers, elicited by high order cognitive functions or disorders, remain to be addressed. This brings to a fundamental question: Is it possible to describe EIB kinetics as a function of complex cognitive manipulations? Along this path, this immediately follows: How do functional and neurochemical information streams contribute to describing brain dynamics and which are the temporal timescales of this relationship?

To address these questions, we propose a novel experimental design by integrating functional neuroimaging and single voxel brain spectroscopy. Using sliding-window and visibility graph analysis, we measure changes in the equilibrium of neurotransmitters across mental workload levels (Figure 1) as a function of time.

### Using spectroscopy kinetics to infer about brain function

Previous 3T fMRS studies have demonstrated that in-vivo concentrations of glutamate^28,29^ and GABA^30^ in the left DLPFC are related to a repeated WM task load and performance, although inconsistencies arise due to variations in experimental protocols. In an attempt to overcome potential attenuators or confounders, starting from a similar paradigm, we use a parametric manipulation of the WM load with the idea of avoiding adaptation effects from simple task repetition and boosting individual differences^31^. In our exploratory approach, we examine EIB by comparing snapshots of integrated metabolic responses to increasing levels of sustained activity. However, the static method hinders the detection of metabolic differences across sessions: we observe no significant changes in mean neurometabolite concentrations from resting-state to the 2-Back condition, when estimated as a static average across each cognitive condition. This observation underscores the transient nature of the phenomenon and highlights the critical importance of incorporating a temporal perspective in our analysis to effectively investigate changes in the balancing of chemical signaling. By adopting this dynamic approach, we were able to uncover more nuanced changes. Specifically, we found that EIB kinetics increased when comparing the 2-Back condition to the other sessions. This finding is aligned with the idea that increasing the WM proficiency parametrically (as seen by behavioural accuracy results of 1-back task ∼98%) leads to a sudden rise in the excitatory neural spiking activity of the prefrontal cortex^32–35^, which is characterized by high glucose demand and oxidative metabolism^16,36^. Moreover, by looking separately at neurometabolites, we found a concomitant increase trend in Glx, thus pointing to a general up-regulation of the glutamatergic cycle during the attentive process or trends to decreased inhibition during learning/repeated stimulation^37–39^. Consistently with previous studies^30,40^, we observed inhibitory kinetics changes during the early periods of the 2-Back task as suggested by recent models of rapid shifts^16^.

While many studies indicate that glutamate levels typically increase in response to cognitive demands, reflecting enhanced but not sustained metabolic activity and excitatory neurotransmission^29,41^, this finding is not universal; some research reports no significant changes in glutamate levels^30^. Conversely, GABA concentrations demonstrate a biphasic temporal profile: an initial increase at task onset followed by a gradual decline in subsequent task runs^30^. This dynamic GABA pattern, in conjunction with glutamate fluctuations, suggests a coordinated metabolic response involving active GABA-glutamate-glutamine cycling throughout WM engagement. Furthermore, the aforementioned interplay is supported by the evidence that effective WM function relies on a well-documented balance between excitatory and inhibitory processes. Lower glutamate levels are associated with improved task performance^29^, suggesting that efficient excitatory neurotransmission, rather than excessive activation, may be optimal. GABA dynamics also contribute significantly: both dynamic changes in GABA+ levels throughout task engagement^30^ and higher baseline GABA concentrations^40^ correlate with enhanced WM capabilities. Moreover, the balance between inhibitory and excitatory neurotransmitters, as reflected by a lower GABA/Glutamate ratio, is linked to superior WM performance^42^. Collectively, these findings highlight the delicate equilibrium required between excitatory and inhibitory processes for optimal cognitive function, emphasizing the importance of nuanced neurometabolic regulation in supporting WM processes.

### Insights into the timescale of neurometabolic and neurovascular coupling

Drawing upon the previously established conclusions, we explored the relationships between dynamic functional connectivity and EIB kinetics, as illustrated in Figure 3. Our findings reveal that cognitive load influences the association between the temporal characteristics of FPN executive functioning and the kinetics of the balance between glutamate (Glx) and GABA+. These results provide a more integrated and time-resolved description of the positive associations observed between neurochemical concentrations and hemodynamic response in both perceptual^21,34^ and highly demanding cognitive tasks^29^. In what follows, we will provide a more comprehensive understanding of how the brain adapts to varying cognitive loads by integrating these two multi-timescale dynamics.

Higher temporal stationarity of the executive control network is thought to be beneficial in sustained attentive and WM processes^43,44^ if characterized by dynamic reconfiguration and load-dependent modulation. Indeed, we probe that temporal stationarity of prefrontal executive networks is associated with EIB kinetics and can differentiate changes in WM load conditions. Temporal features of FPN not only are parametrically stabilizing as a function of load condition (i.e., toughest cognitive performance) but are also highly aligned with EIB kinetics. The signature patterns of association between EIB kinetics and temporal features of FPN probe a positive association between EIB’s kinetics and functional network stability in the highest cognitive load condition (e.g. 2-Back), whereas a reversed pattern is observed for attentive-only task (e.g. 0-Back). Consequently, for high cognitive load conditions where FPN temporal properties are balanced, the EIB curve parametrically changes towards a kinetics characterized by increased temporal gradient and reduced time-invariant evolution. As EIB kinetics increase over time, the excitation counterpart of the ratio (i.e., Glx) is presumably boosted relative to the inhibitory component, suggesting that metabolic characteristics of the DLPFC^18,41^ may be primarily related to glutamatergic neurons.

While our findings provide insights into the relationship between FPN dynamics and EIB kinetics, it’s important to note that neurochemical-dependent changes in brain metabolite spectra remain a subject of debate. Different models offer varying interpretations of these changes. For instance, Lea-Carnall et al. (2023)^16^ developed a computational model simulating MRS-detected shifts in glutamate and GABA, attributing these to rapid exchanges between vesicular and cytosolic compartments that stabilize within about 5 seconds after activity changes. In contrast, Mangia et al. (2009)^17^ propose that glutamate changes in fMRS are due to slower metabolic processes, linking fluctuations to energy metabolism rather than immediate neurotransmitter cycling. These differing interpretations highlight the complexity of understanding the physiological temporal dynamics of neural signalling and its associated vascular response, which operate across time windows that differ by orders of magnitude. While the effects of vascular changes on MR signals are well-characterized, a coherent description of neural signalling dynamics via MRS, and their temporal interplay, remains to be fully established.

Finally, we investigated the timescale of the metabolite and hemodynamic responses in executive network dynamics. In this study, the reported increase in stationarity of the FPN network is further clarified by the dynamics of the persistence probability curve, which shows a sustained increase following the third frame during the most cognitively demanding task. In temporal terms, this corresponds to approximately 24 seconds of WM activity required to detect an increased probability of network persistence. In the context of EIB, similar timescale considerations apply. EIB changes are detectable as early as 12 seconds after baseline, reaching a steady state towards the end of the observation period, around 8 minutes. These findings are consistent with previous studies that propose the coexistence of multiple timescales of the metabolic response: an initial excitatory phase (lasting minutes^21^), characterized by a rapid rise in signal, followed by a slower phase^17,26^, where inhibition, on the order of tens of minutes, establishes a plateau. Moreover, the gradual emergence of the inhibitory component may be undetectable in static EIB analyses, as whole-acquisition averaging obscures the rapid Glx kinetics, which decay within seconds, while favoring the slower component persisting throughout the entire observation window.

The temporal dynamics of the FPN during sustained WM load appear to be tightly coupled to this specific network configuration. The other CAPs obtained from the dynamic connectivity analysis, although including large portions of the DLPFC, do not exhibit persistence features comparable to those of the FPN, which are correlated with the kinetics of EIB. This suggests that the regulation of EIB in the DLPFC and the involvement of the network associated with WM may be closely related. This mutual interaction seems to occur on similar timescales during the initial phase (with both curves showing the aforementioned rapid changes in the first frames, within 12-24 seconds from baseline), and remains elevated throughout the entire measurement window. The mechanism that supports both the elevated persistence on one side and EIB on the other may be due to a redefinition of the baseline equilibrium in EIB, driven by the cited progressive contribution of the inhibitory component, which limits the excitatory signalling likely stimulated by the energy demand required to support WM.

However, this work comes with some limitations. Firstly, the target voxel location, the DLPFC, was proximate to the skull, potentially causing deteriorated spectra given the presence of lipid artefacts and partial volume effects. After quality check, we had to exclude half of the initial sample size, to ensure data adequateness^45–48^ potentially impacting the accuracy and generalizability of our results. Regarding spectral neurometabolites fitting, we only reported composite measures for Glx (i.e., Glu+Gln), given inconclusiveness and low reliability of linear combination modeling approaches^23^. Additionally, a control voxel in a brain region unrelated to the task was omitted due to acquisition protocol time constraints. Instead, we inserted a control condition for the task (i.e., 0-back) and a resting-state session to set a subject-specific baseline, which was later used to create normalized concentration changes of neuro- metabolites across cognitive load conditions. This helps reduce the inter-subject variability of baseline metabolite concentration levels, which can be influenced by various confounds^49^. In order to overcome the issue of brain function localization, further studies should improve fMRS measurement to enable whole-brain fitting of neurometabolites through usage of advanced techniques, such as MRSI^50,51^. Future efforts should be directed towards improving the temporal scale of the reported measurements by including proxy measurement of EIB (e.g., Hurst exponent, intrinsic timescale) derived from other neuroimaging approaches such as EEG with higher temporal resolution^12,14,15^.

In conclusion, using a peculiar combination of fMRS and fMRI, along with time-resolved analyses, we provide evidence of integrated temporal properties of executive networks and EIB kinetics in healthy volunteers. Understanding the temporal dimension through which metabolism and vascular response interact to drive *ad-hoc* reconfiguration of brain networks for information processing is crucial to possibly reframe cognitive disorders in terms of disrupted synchronicity or impaired coupling of neural signals, where present.

## Materials and Methods

### Participants

A total of 36 healthy adult volunteers (17 females) were recruited in this study and a subset of them underwent the whole fMRI-fMRS experiment (N=24, 11 females). The study was approved by the Ethical Committee of University of Trento. Demographic information of the sample is displayed in Table 1, together with potential confounding variables for the GABA-edited MRS^55^. Exclusion criteria encompassed: neurological or psychiatric disorders; current psychotropic medication use; large nicotine consumption (>1 cigarette/day); large alcohol intake (>14 units/week or >3 days/week) and migraine with aura. Participants were required to abstain from caffeine intake 12 hours pre-scan. Given the novelty of the research line, power analysis was based on previous research^44^ and conducted using GPower (v3.1.9.7). The software indicated that a sample size of 22 subjects was adequate to detect a small to moderate within-subject main effect (f ≥ 0.20) across 3 tasks (see Experimental Design) at the recommended power (0.80) and α = 0.05 for autocorrelated measures (r = 0.75). Thus, a sufficient number of participants were included in both the full sample and the fMRS subgroup.

**Table 1:** Demographic information of the healthy volunteer groups that underwent the parametric working memory task during functional MRI or during the combined functional MRI and MRS protocol (Figure 4). Details are reported using standard guidelines for MEGA-PRESS usage (Peek et al., 2023)

| Sample | fMRI | fMRI-fMRS |
| --- | --- | --- |
| **Sample size** | 36 | 24 |
| **Age (years)** | 25.6±3.2 | 25.0±3.1 |
| **Gender** | 19 M, 17 F | 13 M, 11 F |
| **Education (years)** | 18.8±2.5 | 18.8±2.5 |
| **Handedness** | 5 left-handed | 5 left-handed |
| **Menstrual cycle** | N.A. | Menstruations (1-5 days): 1<br>Proliferation phase (6-14 days): 3<br>Ovulatory phase (14-15 days): 1<br>Initial secretion (16-23 days): 2<br>Final secretion (14-28 days): 3<br>Amenorrhea: 1 |
| **Nicotine** | N.A. | 10 nicotine smokers |

### Experimental design

The experimental procedure aimed at non-invasive in-vivo mapping of both EIB derived from single-voxel MRS and brain function derived from BOLD fMRI while manipulating the cognitive load level in healthy volunteers (schematically represented in Figure 4). The design consisted of four different MRI sessions, each of them interleaving an fMRI and an fMRS acquisition. The first session was designed as a baseline condition, called resting-state (TAfMRI-fMRS∼14 min), where the participants were instructed to lay still in the scanner, think about nothing in particular and fixate the crosshair displayed at the centre of the screen. The following three fMRI/fMRS sessions were acquired during a visual WM task. In particular, in order to parametrically increase the level of WM load, participants were asked to perform a 0-Back, a 1-Back and a 2-Back task in the first, second and third session, respectively. The order of the tasks was not shuffled across subjects given previous studies about the timing of the expected metabolic response in the fMRS session^56^ and to ease inter-individual difference analyses^25^. Each session consisted of a block design, with 4 blocks running during the fMRI and 4 blocks running during the fMRS acquisition. To control for the effect of the functional specialization of the brain areas involved in the WM system^57^, the experimental stimuli consisted of letters and digits, such that the stimulus type was kept constant throughout the 4 blocks running during the same acquisition (i.e., fMRI or fMRS) and varied in the following 4 blocks (except for one session in which, due to an unexpected technical problem, a participant was presented with letters as stimuli during both the fMRI and the fMRS acquisition blocks). The assignment of stimulus type (i.e., letters or digits) to acquisition type (i.e., fMRI or fMRS) was pseudo-randomized across sessions and participants.

**Figure 4:**
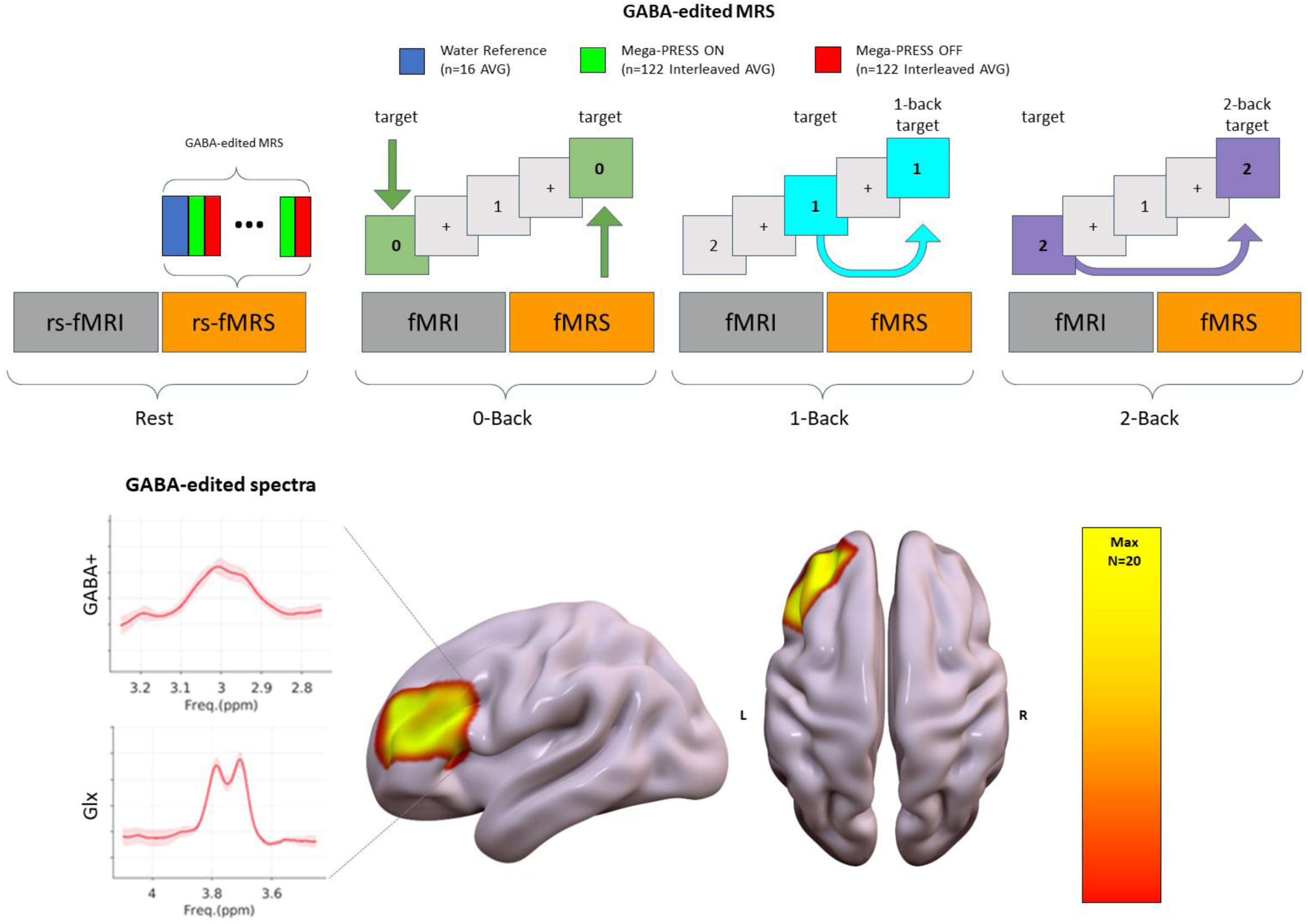
Exploring in-vivo EIB kinetics as a function of cognitive load. First row displays the experimental design consisting of four different sessions (session_1_=Resting-state, session_2_=0-Back, session_3_=1-Back, session_4_=2-Back), each one with interleaved fMRI-fMRS acquisitions lasting approximately 14 min in total per session. For each task session, visual stimuli were randomly selected as digits or letters. Before the beginning of each GABA-edited fMRS acquisition, a water reference sequence was acquired by disabling water suppression. Second row shows the probabilistic anatomical overlap of the MRS voxel placement across subjects in the left dorso-lateral prefrontal cortex (DLPFC) (Max: corresponds to 20 subjects overlapping with the total sample of 24 subjects) together with example mean spectra from the resting-state acquisition across all subject to identify GABA+ and Glx peaks. Abbreviations: MRS (Magnetic Resonance Spectroscopy), AVG (averages), rs (resting-state).

Regardless of the task and stimulus type, each block consisted of 44 trials and each trial consisted of a 2-second stimulus presentation followed by a 0.5-second crosshair presentation. In addition, in the only case of the 0-Back task, participants were instructed to press a button every time they saw a stimulus identical to the target, displayed at the beginning of each block for 2 seconds. Instead, in the 1-Back and 2-Back tasks, participants were instructed to press a button every time they identified a stimulus that matched the one presented 1 and 2 trials earlier, respectively. Irrespective of task and stimulus type, each block comprised 9 congruent trials, that is trials in which participants were expected to respond according to the task instructions.

### Behavioural Analysis

Behavioural data were collected during the three task-based sessions to control for the expected effect on the cognitive load throughout the whole acquisition. Reaction time (RT, response time to congruent stimuli in ms), total accuracy (percentage of correct responses for all trials), hit rates (proportion of recognized congruent trials) and false alarms (proportion of incongruent trials recognized as congruent) were calculated for each subject across different sessions. Moreover, a sensitivity measure^58^ (*d’*) was calculated based on the standardized difference between the false alarms and hit rates distributions for each subject, after adjusting perfect scores by employing a log-linear approach^59^.

### MR Acquisition

Data were acquired with a 3T clinical MR scanner (MAGNETOM Prisma, Siemens Healthcare, Erlangen, Germany), equipped with a 64-channel head-neck RF receive coil and parallel transmit RF (operating system VE11C). The overall MR acquisition protocol obtain after optimization (see Supplementary materials) included:

- Structural 3D T1-weighted multi-echo MPRAGE^60^ (TR=2.5s, TEs=[1.69, 3.55, 5.41, 7,27 ms], TI=1100 ms, 1mm-isotropic voxels)
- Four sessions of functional BOLD 2D EPI (TR=2s, TE=28 ms, 3mm-isotropic voxels, full brain coverage, AC-PC orientation)
- Double-echo gradient echo sequence was acquired to later control geometric distortions of fMRI data (TR 682ms, TE1/TE2 4.2/7.4ms, 3mm-isotropic voxels).
- Four sessions of MRS water reference^61–63^ (MEGA-PRESS; TR/TE=2000/68 ms, editing pulses=1.9/7.5 ppm, editing pulses bandwidth=60 Hz, acquisition bandwidth=2000 Hz, cubic voxel size=3 cm, spectral points=2048, deactivated VAPOR water suppression, 32 averages, TA=2:08 min)
- Four sessions of edited functional MRS^64^ (MEGA-PRESS; TE/TR= 68/2000 ms, brain-optimized automated shimming routine, editing pulses=1.9/7.5 ppm, editing pulses bandwidth=60 Hz, acquisition bandwidth=2000 Hz, cubic voxel size=3 cm, spectral points=2048, VAPOR water suppression, resting-state=192 averages, task=224 averages, TA_REST_=6: 50 min TA_TASK_=7:40 min).

The MRS package was developed by Edward J. Auerbach and Małgorzata Marjańska and provided by the University of Minnesota under a C2P agreement. MRS data were collected using automated voxel placement (described in the Supplementary Materials).

### MRS Preprocessing

MRS data were processed with Gannet v3.2.0 (https://github.com/markmikkelsen/Gannet.git) on Matlab version 9.5.0, a Matlab based analysis tool specifically developed for the analysis of GABA-edited MRS data^65^.

In order to process edited MRS, pre-processing steps included^48^: (i) gross head motion correction; and (ii) phase and frequency drift correction with a spectral registration algorithm^66^. After these corrections, the Fourier transform is computed to perform the frequency spectra subtraction between the ON and OFF spectra, which are aligned and averaged^65^. Once the pre-processing is performed, the neurometabolite spectra peaks are extracted by peak integration fitting targeting GABA+ and Glx (compound metabolite of Glutamate+Glutamine) in the difference spectrum.

The fitted spectra were then corrected by considering the proportion of different tissue types in the voxel, namely gray matter (GM), white matter (WM) and CSF. To do this, the single voxel was co-registered to the subject anatomical image with SPM12 and then segmented in different tissue types. Concentrations of metabolites of interest were quantified relative to the internal water reference. Therefore, the unitless signal intensity values are converted into absolute concentrations with a computation weighting for the percentage of the tissue type involved^67^ enabling comparison between different subjects.

Finally, the macromolecules (MM) contribution was considered by using the Gannet model^48^, which assumes that 45% of the area of GABA+ peak is attributable to MM contribution^65^.

### MRS quality control

Data exclusion criteria were established for the processing methods (see Supplementary materials), taking into account the type of fitting procedure. Quality metrics include: GABA Signal-to-noise ratio (SNR) and fit error, Glx SNR and fit error^51,55^. After computing the fMRS quality metrics for each subject, a distribution across the group was obtained. From this, a z-score of each metric was calculated for each subject relative to its corresponding metric distribution. Data were excluded if the absolute value of the z-score was higher or equal to one in at least two quality metrics or in at least one session for the same metric. Data outliers were calculated during the first analysis described in the following paragraph and removed from further analysis^51^.

### Neurometabolites estimation: static and dynamic

In order to evaluate neurometabolite concentration changes in terms of EIB across sessions with different WM cognitive loads, a two-fold analysis was performed. Firstly, the overall average concentration was averaged across each full MRS acquisition session, separately for resting state (192 averages, 96 ON and 96 OFF), 0-, 1- and 2-Back (224 averages, 112 ON and 112 OFF) conditions. The averages for each MRS session were processed together in Gannet (GABAGlx model). The mean concentration of the neurometabolites during the first session (i.e. resting-state) was later used as subject-specific baseline for further analysis.

Secondly, to evaluate kinetics over different sessions, we computed sliding window averages. Briefly, 60 averages (60 ON and 60 OFF) were taken as window length (4 minutes), which ensured adequate SNR of GABA+ estimates. This window was then shifted in steps of 3 averages (12 seconds), resulting in 18 dynamic frames spanning cognitive conditions beside the resting-state, where they correspond to 12 frames. A small sub-set of averages was discarded in the resting-condition in order to maintain the same frame dimensions over the different sessions. As a way of estimating individual fluctuations of EIB and its associated metabolites over time, curves were normalized over steps using the baseline mean value estimated in the first frame of the corresponding cognitive condition. This step is introduced to control for carryover effects and baseline shifts from previous sessions. Given the lower stability of the fitting due to the reduced number of averages, a median interpolation of outliers was applied. With the aim of gaining insights about the temporal changes of neurometabolites, temporal properties of EIB, Glx and GABA+ changes across sessions were calculated. The limited number of time points available makes the estimation of conventional kinetics properties (e.g. time to peak, slope and zero-crossing) more sensitive to transients and local instabilities. To better characterize the temporal features of the EIB we opted for a graph-theoretical approach. The visibility graph, described in Lacasa et al., 2008^68^, translates time series into graphs that inherit the temporal properties of the original data. We then considered: 1) the average out-degree of the associated time-directed graph as a surrogate metric for the slope (lower average out-degree indicates time-dependent changes of EIB); 2) the KLD - the distance between in- and out-degree distribution - as a proxy for stationarity of the EIB curve^69^ (lower values are associated with time-invariant, or stationary, processes).

### Functional MRI pre-processing

The pre-processing of resting-state and n-back fMRI conditions was implemented using an FSL-based automated pipeline (https://github.com/tambalostefano/lnifmri_prep) that included common steps: (1) slice timing correction; (2) T1-weighted image tissue segmentation; (3) geometric distortion and head motion correction; (4) co-registration of the T1-weighted image to the time-series. For the resting state fMRI sessions, further steps included: (5) regression from the time-series of the 6 head motion parameters, white matter and CSF signals; (6) band- pass temporal filtering [0.01-0.10 Hz]. Both resting-state and task-based fMRI were warped into common space: (7) normalization to standard MNI template space; and (8) spatial smoothing 6 mm FWHM Gaussian kernel size.

### Functional co-activation patterns

To estimate brain fMRI recurrent states associated with in-vivo EIB and modulated by cognitive load levels, we applied a co-activation patterns (CAPs) framework for each fMRI session (https://github.com/MIPLabCH/TbCAPs). In our case, CAPs were obtained by defining a seed in the left DLPFC^57^ (10 mm radius) matching the MNI coordinates of the voxel placement used in the fMRS scans. This allowed identifying the seed BOLD time course.

Then, the timepoints where the seed BOLD activation exceeded a z-score of 1 were selected for further analysis^70^. The fMRI frames of the selected timepoints were fed into a k-means clustering procedure to reconstruct temporal assignments of the spatial patterns. To estimate the best fitting k, a consensus clustering was performed^71^. Lastly, all frames assigned to the same state were averaged to reconstruct the corresponding CAP. Therefore, absolute z-score co-activations patterns were extracted with respect to the left DLPFC seed, for limited periods of the time-series, with clustering aiming at detecting the re-occurring fronto-parietal spatial patterns.

Temporal metrics of each CAP (including the non-active state, CAP_0_) were computed for the different fMRI cognitive conditions, by concatenating subjects from all sessions: (i) in-degree: how likely a CAP is visited from any other one, (ii) out-degree: how likely a CAP is exited towards any other one, (iii) resilience: the likelihood to remain in the same configuration throughout the time-series, (iv) occurrences: how much a CAP re-occurs over the time-series, (v) betweenness centrality: how important a CAP is regarding the shortest paths between other CAP pairs^70^.

### Statistical analysis

For the evaluation of the behavioural performance, the statistical testing was carried out for both fMRI blocks in the whole sample and fMRS blocks in the fMRS subgroup using R (v4.3.1). The Shapiro-Wilk test for normality was used to assess the distributions of the variables for accuracy, *d’* and RT for each task. After removing outliers (i.e., values 1.5 times the interquartile range greater than the third quartile or 1.5 times the interquartile range less than the first quartile), mean accuracy and *d’* were compared using the Friedman test as a non-parametric alternative to the repeated measures ANOVA, and the Wilcoxon signed-rank test for performing post-hoc pairwise comparisons (Bonferroni correction). For the RT analysis, a generalized linear model (GLM) was fitted using the Gamma distribution and comprised RTs as dependent variable, five fixed-effect factors (the three sessions, the four blocks for each session, the number of responses provided by each participant during each session, participants’ gender and age), a random intercept by participant and a random slope by session. Tukey correction was then applied to estimated marginal means for multiple comparisons. The significance level alpha was set to 0.05.

In addition, the same GLM was employed to test for the link between behavioural performance and the static concentration of neurometabolites, their kinetic characteristics, and the same attributes of FPN. Since the static concentrations were already controlled for age and gender, the only fixed-effect component included in the model was the number of sessions together with an interaction term for the variable of interest across sessions. Furthermore, this methodology was employed to investigate the correlations between static concentrations of neurometabolites and the temporal properties of the fronto-parietal network (FPN). Delta GABA+ levels were derived by subtracting the resting state concentration from the measured GABA+ levels to account for baseline fluctuations and to better isolate the dynamic changes in the neurometabolite associated with FPN activity. For the evaluation of cognitive load condition effects on static and dynamic fMRS metrics, the neurometabolite concentrations on each condition were first normalized in relation to the baseline values. For the static fMRS estimates, the whole resting-state value was considered. For the dynamic fMRS estimates, the first frame of each condition was used to control for carry-over effects from previous fMRI manipulations.

For the static fMRS analysis, a Kruskal-Wallis test was performed on the normalized Glx and GABA+ concentrations together with EIB ratio across different sessions (i.e. *Load* of: Rest, 0- Back, 1-Back and 2-Back), which were considered as within factors. Post-hoc paired comparisons were employed to understand the directionality of the effects.

For the dynamic fMRS analysis, the area under the curve (AUC, expressing the metabolite modulation over time) was calculated for each dynamic time-series and statistically tested with a Kruskal-Wallis test on EIB ratio, GABA+ and Glx concentrations across different sessions (Rest, 0-Back, 1-Back and 2-Back) as within factor (i.e. *Load*). No further covariates were included given the inter-individual normalization step. Post-hoc comparisons were employed to understand the directionality of the effects.

For the dynamic fMRI analysis, a Kruskal-Wallis test was performed on the temporal features of FPN-CAP across different sessions (i.e. *Load* of: Rest, 0-Back, 1-Back and 2-Back), which were considered as within factors. Post-hoc paired comparisons were employed to understand the directionality of the effects.

For the fMRI-derived measures, to better understand the relationships between the temporal features of the FPN-CAP and neurometabolites kinetics across different sessions, we applied partial least squares correlation^28^ (PLSC).

The PLSC methodology is a multivariate statistical method that finds the latent variables, or mutually orthogonal, weighted linear combinations of the original variables in the two datasets that have the highest degree of correlation with one another. In the current analysis, one dataset represents the temporal properties of the FPN-CAP (i.e., X_n×t_) with n=12*5 rows as the sample size across sessions where the network was detected [rest, 0-Back 1-Back, 2-Back] and t =5 columns as main temporal properties for the FPN-CAP of interest (i.e. in-degree, out-degree, occurrences, betweenness centrality and resilience). The other dataset the temporal properties of EIB kinetics (i.e., y_n×m_) with n=12*5 rows as the sample size across sessions where EIB was calculated [Rest, 0-Back, 1-Back, 2-Back] and m=2 columns as main temporal properties for metabolic counterpart of EIB kinetics from visibility graph analysis (i.e. the average out-degree and KLD).

Both data matrices were normalized column-wise (i.e., z-scored) in order to identify the latent variables. The correlation matrix R=X’Y was then subjected to the following singular value decomposition: R=X’Y=USV’ where S_m×m_ is the diagonal matrix of singular values and U_t×m_ and V_m×m_ are the orthonormal matrices of the left and right singular vectors, respectively. A latent variable corresponds to each column in the **U** and **V** matrices. Each element of the diagonal of **S** is the corresponding singular value. The temporal FPN-CAP features’ and temporal EIB features’ relative contributions to latent variables are shown by the left and right singular vectors, U and V, respectively.

Positively weighted temporal FPN-CAP features correlate with positively weighted temporal EIB features, whereas negatively weighted temporal FPN-CAP and temporal EIB kinetics correlate with each other. Brain scores show how much each area of the brain displays the weighted patterns found by latent variables that can be estimated using singular vectors. The computation of brain scores for temporal CAP and temporal EIB kinetics characteristics involves projecting the initial data onto the weights that are determined from PLS, specifically **U** and **V**, obtaining:

- Brain scores for temporal FPN-CAP features = **XU**
- Brain scores for temporal EIB features= **YV**

The Pearson correlation coefficient between the original data matrices and the relevant brain scores is then used to calculate loadings for temporal FPN-CAP features and temporal EIB kinetics features. The correlation coefficients between the initial temporal FPN-CAP characteristics vectors and the PLS-derived brain scores for temporal FPN-CAP features, for instance, are known as temporal FPN-CAP features loadings. Using 10000 permutation tests, the statistical significance of latent variables was evaluated. The original data was randomized using spatial autocorrelation-preserving nulls. Every permutation was subjected to the PLS analysis once more, producing a null distribution of singular values. After that, the original singular values’ significance was evaluated in comparison to the permuted null distributions (Figure 3A). Using bootstrap resampling, which involves randomly resampling rows of the original data matrices **X** and **Y** 10000 times with replacement, the dependability of PLS loadings was assessed. Next, for every resampled data set, the PLS analysis was performed once again to provide a sampling distribution for every temporal FPN-CAP feature and temporal EIB kinetics feature (i.e., 10000 bootstrap-resampled loadings). We next utilize the bootstrap-resampled loading distributions to determine the loadings’ 95% confidence intervals (e.g., see Figure 3).

Finally, a sliding-window analysis was conducted on FPN-CAP expression indices to further explore the temporal scale similarities with EIB kinetics temporal features. Identical window features were applied (based on the TR of 2s): 120 timepoints were taken as window length (4 minutes) with shifts in steps of 6 timepoints (12 seconds), resulting in 15 dynamic frames spanning cognitive conditions including the resting-state. To investigate the temporal curve, we report the percentage probability of persistence of the given CAP in subsequent timepoints across the window intervals. Afterwards the area under the curve (AUC) was calculated for each normalized persistence probabilities curve and points were statistically tested with a Kruskal-Wallis test across different sessions (Rest, 0-Back, 1-Back and 2-Back) as within factor (i.e. *Load*). Post-hoc comparisons were employed to understand the directionality of the effects.

## Acknowledgements

We thank Malgorzata Marjańska and Edward J Auerbach, University of Minnesota, for making their MEGA-PRESS sequence available. We would like to express our appreciation to Prof. Micheal Lombardo and Dr. Ludovico Coletta for fruitful discussions about the neuroscientific relevance of these results. We also thank Dr. Stefano Renzetti for the support on the choice of statistical analysis.

## Funding

ISMRM Exchange Award 2021-2022 “Investigating in-vivo human brain dynamic connectivity with fast fMRI”, Dipartimento di Eccellenza project 2018-2022 (Italian Ministry of Education, University and Research).

## Author contributions

**Francesca Saviola:** Writing - original draft, Formal analysis, Project administration, Visualization, Software, Investigation, Writing - review & editing, Conceptualization. **Stefano Tambalo:** Formal analysis, Visualization, Software, Writing - review & editing. **Laura Beghini:** Formal analysis, Visualization, Software, Writing - review & editing. **Asia Ferrari:** Formal analysis, Project administration, Writing - review & editing. **Barbara Cassone:** Formal analysis, Writing - review & editing. **Dimitri Van De Ville:** Writing - review & editing, Investigation. **Jorge Jovicich:** Supervision, Writing - review & editing, Investigation, Supervision, Conceptualization.

## Competing interests

Authors declare that they have no competing interests.

## Data and materials availability

All code, sample data and materials used in the analyses are available (https://doi.org/10.5281/zenodo.14196482) to any researcher for purposes of reproducing or extending the analyses including the v1.0.0 of the Github repository (https://github.com/CIMeC-MRI-Lab/Brain-metabolism-and-connectivity-dynamics). Raw data materials can be requested to the corresponding author through materials transfer agreements (MTAs).

## Notes

### Competing Interest Statement

The authors have declared no competing interest.

### Summary of Updates

This version of the manuscript has been revised for article submission to peer-reviewed journals.

