## Supplementary Materials for "Multiscale excitation-inhibition balance dynamics: integrating metabolite kinetics with time-varying executive networks"

#### Supplementary Materials and Methods

1.1 Protocol optimization
1.2 Edited fMRS acquisition and quality assessment
1.3 Edited fMRS Preprocessing

#### Supplementary Results

2.1 Behavioural performance tests
2.2 Co-activation patterns analysis
2.3 Static edited fMRS
2.4 Dynamic edited fMRS

References

### Supplementary Materials and Methods

#### 1.1 Protocol optimization

The acquisition protocol was optimized by running test scans both on a SPECTRE (Spectroscopy Reference: https://goldstandardphantoms.com/products/spectre/) phantom from Gold Standard Phantoms and in vivo. The optimization aimed to fine-tune the interleaved fMRS-fMRI experimental design to meet this study's objectives, and to obtain fMRS data that satisfy the quality standards[1,2]. In particular the optimization focused on establishing: (i) a consistent voxel placement procedure across subjects; (ii) tuning the acquisition TR; (iii) comparing two spectral alignment algorithms; (iv) assessing the water peak field drift and its effect; (v) measuring the metabolites peaks SNR and FWHM to evaluate the spectral quality; (vi) evaluating the fit error as a goodness-of-fit metric; (vii) considering the desired temporal resolution and metabolites SNR, (viii) and defining the optimal duration for the fMRS runs in the interleaved design.

#### 1.2 Edited fMRS acquisition and quality assessment

##### 1.2.1 Automated voxel placement for MRS acquisition

Before starting the MR acquisition and following the description of the overall experiment, participants were positioned in the scanner and instructed to relax and stay still for the entire duration of the scanning session. After acquiring the anatomical image ME-MPRAGE, the root mean square (RMS) reconstructed image was given as input to an Automated Voxel Placement (AVP) suite[3] to avoid operator-biased voxel positioning. The AVP algorithm translates standard coordinates in MNI space into subject space location on the RMS ME-MPRAGE. This allows consistent MRS voxel positioning across different subjects (between-subjects voxel overlap > 94%, within-subject overlap > 97%, see Fig. 1A for a summary) by moving a template voxel defined in MNI to its subject specific version. To probe the modulation of EIB during a WM task, the template voxel was placed in MNI coordinates=[-36, 44, 20] left dorsolateral prefrontal cortex (l-DLPFC)[4]. The computational time to calculate the voxel coordinates in the subject space resulted in about five minutes across participants. We decided to use the AVP procedure because it improves voxel positioning reliability across subjects relative to manual voxel placement: about 70% between-subjects voxel overlap and about 94% within-subject overlap[3]. Previous studies have demonstrated that metabolite levels differed by up to 30% depending on voxel placement, and indeed inconsistent voxel placement between scans is an often-overlooked source of error when acquiring MRS data[3]. Our procedure includes a minimal editing of two scripts included in the AVP suite (Figure S1): (i) AVP create: to design a template voxel centred on the relevant location in the MNI space; and (ii) AVP co-register: to transform the template voxel coordinates from the MNI space to the subject space. To provide more flexibility, the scripts were adjusted to include in the output some of the intermediate steps of the processing; other edits were included to reduce the computational time and a brain-extraction step was included before the linear registration. None of these manipulations altered the main logic and flow of the algorithm. To check consistency in the voxel placement across subjects with AVP we converted the voxel coordinates returned by AVP from the subject space back to the MNI space (with the normalize function in SPM12 (https://www.fil.ion.ucl.ac.uk/spm/software/spm12/). Then we binarized the result and summed voxel images across subjects using FSL[5]. Subsequently, we overlaid the voxels on an MNI template brain image displaying the probabilistic map of the voxel placement (Fig. 1A).

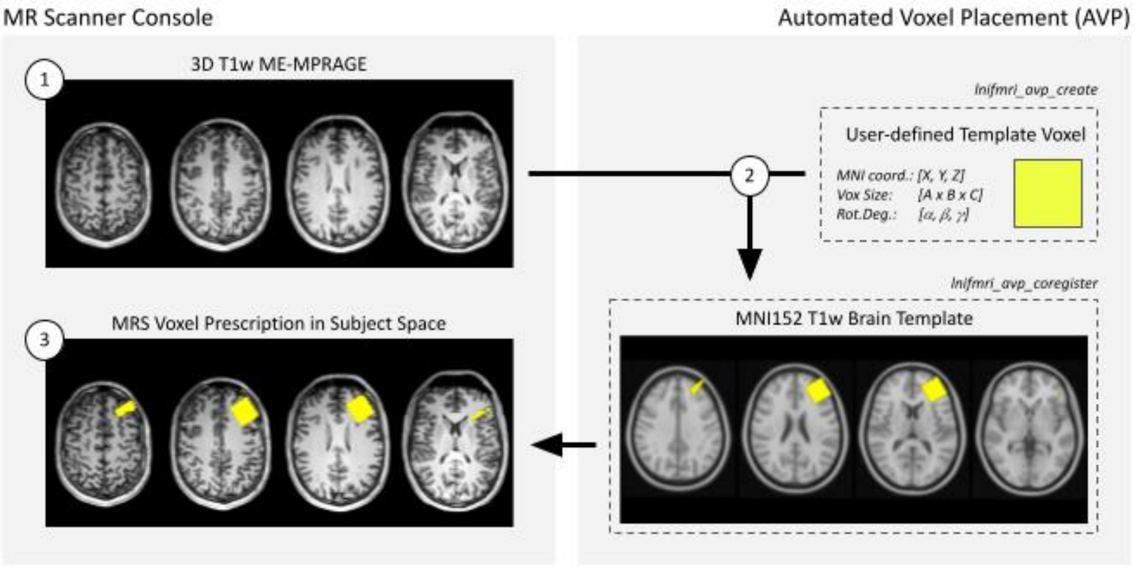


Figure S1. Schematic representation of the automated voxel placement workflow implemented in this work. Root-mean-square reconstruction of high-resolution ME-MPRAGE is exported (step 1) in DICOM format to an external computer running the customized AVP suite (step 2, see main text for details of the customization). Here, the user defined target voxel template is co-registered from MNI space to subject space (step 2). Finally, subject-specific voxel coordinates are returned and entered into the sequence parameters card of the MR scanner (step 3).

##### 1.2.2 Optimization of TR choice for Edited fMRS

Pilot data were collected on three healthy subjects, placing the voxel in the posterior cingulate cortex (PCC). In order to investigate the effect of temporal resolution, the MEGA-PRESS sequence was run twice (TR=1.5 s, spectral points=1024 or TR=2 s, spectral points=2048). The GABA+ and Glx signal to noise ratio (SNR) did not show significant differences across TRs (Figure S2). However, a positive trend was shown for TR=1.5 s (GABA+: 24.9 ± 5.5, Glx: 22.7 ± 3.1) compared to TR=2 s (GABA+: 24.1 ± 3.1, Glx: 26.5 ± 5.8) for an acquisition time of 640s. A TR of 2 s is commonly used for MEGA-PRESS sequences targeting GABA and Glx[6]. We decided to stick with the most common temporal resolution to facilitate results comparison with literature, as the considered alternative did not significantly increase the target metabolites SNR.

##### 1.2.3 Optimization of spectral alignment choice for Edited fMRS

Two spectral alignment methods, Cr alignment and RobustSpectralRegistration (rSpecReg[7]), were compared on the pilot data collected in PCC on three healthy volunteers. The spectral quality was visually improved when using the Robust Spectral Registration Algorithm compared to the Cr alignment (Figure S2, panels A,D). When using the Cr alignment the GABA+ peak full width half maximum (FWHM) (17.7±2.8) Hz was on average smaller compared to the Robust Spectral Registration case (19.6±2.0), independently of the TR (Figure S2 B). This heldfor the Glx peaks as well (Cr-alignment=11.6±2.1; rSpecReg=13.2±1.2) (Figure S2, panel E). On the contrary the GABA+ and Glx SNR (Figures S2 C,F) were higher when using the rSpecReg compared to the Cr algorithm (SNR GABA+ rSpecReg=24.5±6.3; SNR GABA+ Cr-alignment=22.4±7.1; SNR-Glx rSpecReg=24.6 ±7.1; SNR-Glx Cr-alignment=22.4±8.6). Therefore, for our study we adopted the rSpecReg since it leads to bigger spectral peaks (i.e. higher SNR and higher FWHM), outperforming the Cr alignment approach.

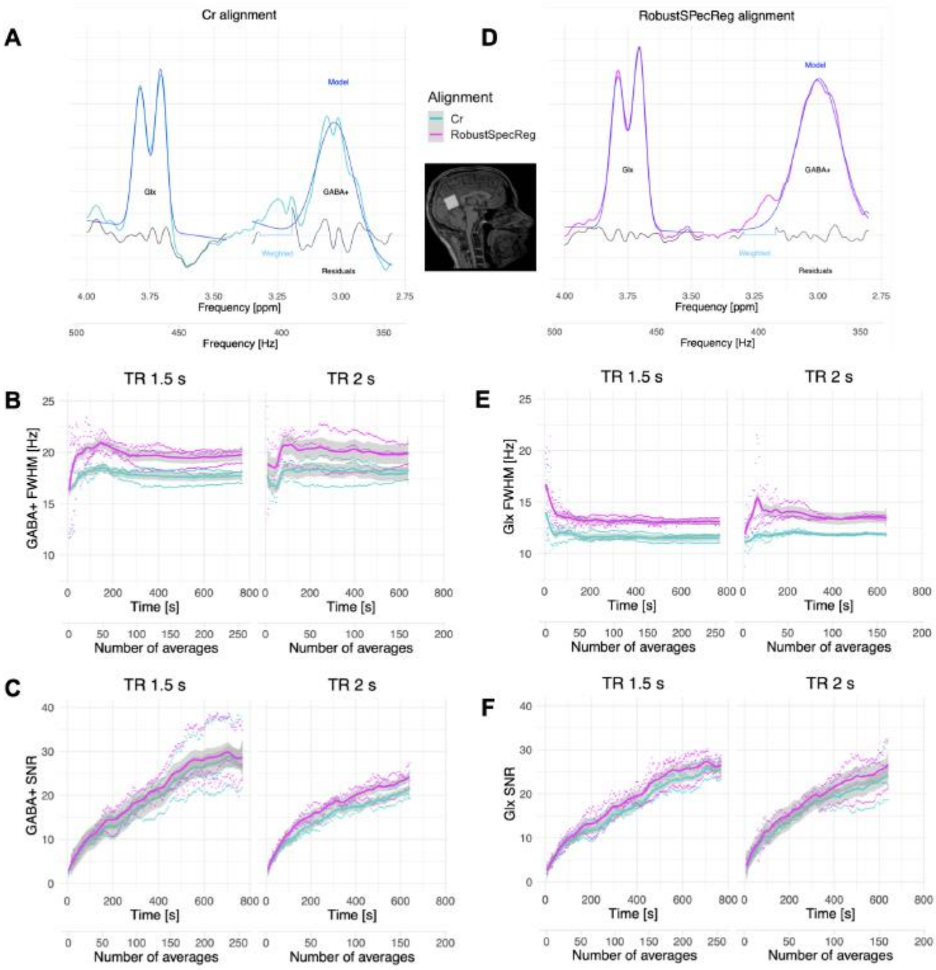


Figure S2: The TRs and alignments methods comparison. In this figure only the data collected in the posterior cingulate cortex (PCC) with both repetition times (TR 1.5 s and 2s) are shown. On panel A and D the spectra collected with TR 2s in a representative subject are shown (with the same y-scale). The Cr alignment and Robust Spectral Registration algorithm were used to obtain the spectra on panel A and D respectively. On panel B and E the GABA+ and Glx full widths half maximum (FWHM) dependence on the acquisition time and number of averages are shown for both TRs and both alignment methods. On panel C and F the GABA+ and Glx signal to noise ratio (SNR) dependence on the acquisition time and number of averages are shown for both TRs and both alignment methods. The solid lines in panels B, E, C, F were obtained averaging on the three test subjects' data.

Following the results on optimization of frequency and phase alignment between transients via rSpecReg, both on the phantom and human pilot experiments, the same method was applied in the pre-processing of static MRS data.

The spectral registration method was reconsidered in the context of time dependent metabolite quantification. Given the lower intrinsic SNR in both GABA and Glx within the sliding window size (each frame consisting of 60 edited spectra), we explored different methods to optimize the spectral alignment. We considered linewidth (FWHM) and SNR as a function of rSpecReg and SpectralRegistration (SpecReg[8]) algorithms, against the simplest case of no alignment. Figure S3 shows the effect of spectral registration on a representative subject. Qualitative improvement of spectral quality is visible especially in the GABA complex, which exhibits narrower peaks compared to the raw edited spectrum. Average values for the two metrics are presented in Table S1, visual comparison of framewise quality metrics for GABA and Glx, is reported in Figure S4. Significant improvement in spectral quality, expressed as reduction of FWHM, were found when using SpecReg both for GABA ($F(2)=79$, $p<<0.01$) and Glx ($F(2)=145$; $p<<0.01$). In terms of SNR, non-significant improvements were observed. Based on these results, we opted for the SpecReg algorithm to control for spectral misalignment.

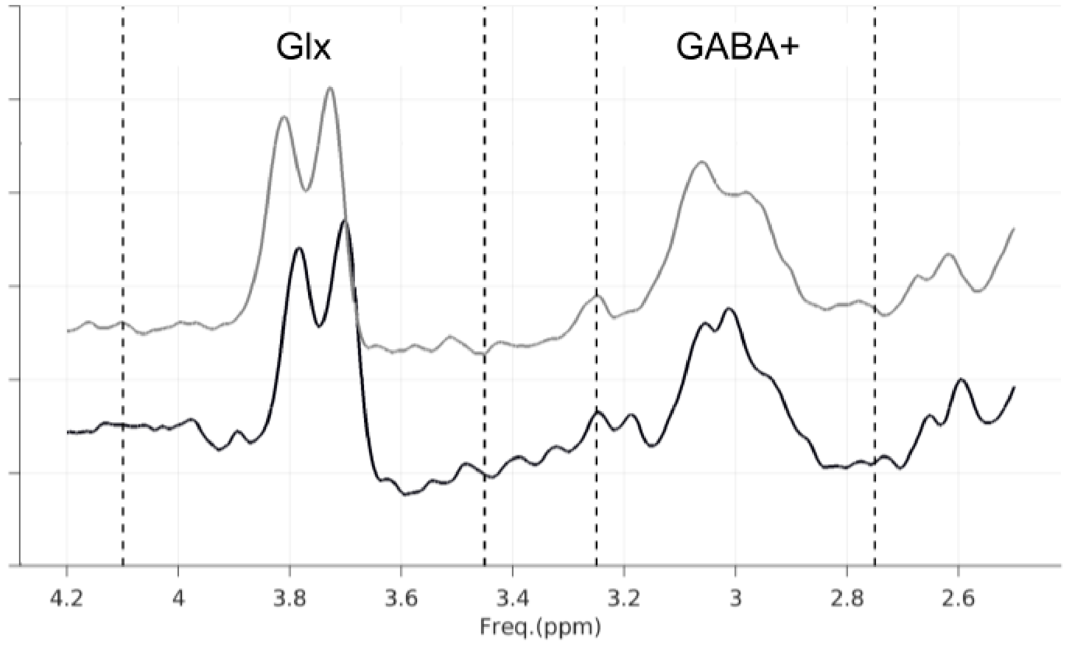


Figure S3: **Effect of spectral registration.** Representative edited spectrum before (gray line) and after (solid black line) spectral alignment from a sample subject. Glx and GABA+ frequency bands (3.45-4.1 and 2.75-3.25ppm, respectively) are marked by dashed lines. Vertical displacement of spectra is introduced for visualization purposes.

| | **SNR** | | | **FWMH** | | |
|---|---|---|---|---|---|---|
| | none | SpecReg | rSpecReg | none | SpecReg | rSpecReg |
| Glx | 18.6 ± 8.2 | 18.3 ± 3.7 | 19.1 ± 5.4 | 15.3 ± 2.5 | ***12.8 ± 1.1 | 16.5 ± 7.0 |
| GABA | ***15.5 ± 2.7 | 17.3 ± 5.0 | 17.2 ± 5.8 | 19.3 ± 1.8 | ***16.6 ± 5.7 | 18.9 ± 5.5 |

Table S1. **Comparative Spectral Quality Metrics produced by different alignment methods.** Data are reported as mean ± standard deviation for Glx and GABA. Asterisks indicate post hoc significant differences from a one-way ANOVA ($p<<0.01$), detailed results are reported in the paragraph. SpecReg: spectral registration (Edden et al., 2014); rSpecReg: robust spectral registration (Mikkelsen et al., 2020).

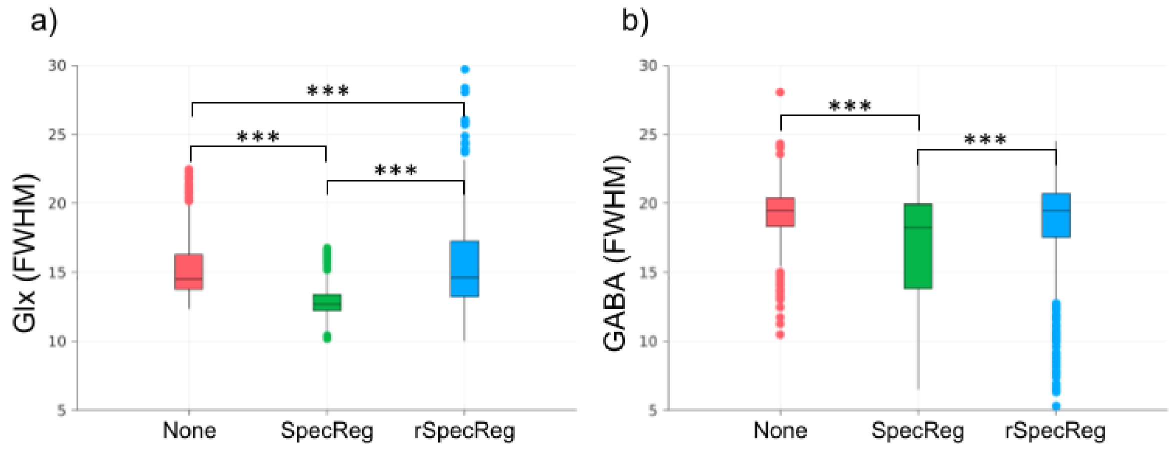


Figure S4. **Comparison of three different spectral alignment methods evaluated in the preprocessing of fMRS data**. A significant reduction in FWHM was obtained both for Glx, panel a), and GABA, panel b), in the case of SpectralRegistration. Asterisks indicates posthoc significant differences from a one-way ANOVA ($p<<0.01$). SpecReg: spectral registration (Edden et al., 2014); rSpecReg: robust spectral registration (Mikkelsen et al., 2020).

##### 1.2.3 Investigating the effect of interleaving fMRI with fMRS

The BOLD EPI sequence is known for causing scanner heating leading to field drifts[9]. These can result in a reduction of the editing efficiency of the MEGA-PRESS sequence, which relies on narrow band frequency-selective pulses. To the aim of investigating the scanner field drifts and heating effects on the MRS spectral quality, the interleaved fMRI-fMRS protocol was run on the phantom. The acquisition was run starting with a cold and non-cold scanner as the initial scanner condition is known to affect the water peak drifts intensity[9].

The water peak frequency drifts and their effect on the GABA SNR are shown in Figure S5. Six interleaved and consecutive sessions were run starting with a cold scanner. In the first session (TR 2s), the drift was about four times higher ((5.93±0.09) Hz/640s) compared to the sixth session ((1.53±0.09) Hz/640s). We noticed that drifts higher than 2.5Hz/640s (as in the first three sessions with TR 2s) cause a deformation in the GABA spectral peaks (Figure S5 C) and a reduction in the SNR (Figure S5 D). Indeed, GABA SNR for the data collected in the first session (7.5±0.5) is almost halved compared to the last session (13.0±1.0).

Three interleaved and consecutive sessions were run starting with a non-cold scanner, with the voxel positioned both in the center of the spherical phantom and near the phantom's surface. The GABA peaks in these two cases did not show visible deformation due to the water drifts (Figure S5 C). The water peak frequency drift (Figure S5 B) was substantially reduced compared to the cold scanner case and the GABA SNR increased. On average higher drifts were measured, when placing the voxel close to the phantom surface ((0.95±0.47) Hz/640s) compared to the centred position ((0.85±0.38) Hz/640s). On average, higher GABA SNR (Figure S5 E) was measured, when placing the voxel close to the phantom surface (64.6±7.2) compared to the centred position (17.2±3.0). The drift measurements obtained with cold and non-cold scanner are in the expected range[9].

The GABA editing RF pulses acts on a 60 Hz frequency bandwidth and we can assume them to give rise to a Gaussian excitation in the frequency domain[1]. Therefore, a 2% frequency (±

1.3 Hz) shift in B0 is not expected to affect RF editing efficiency for GABA measurements. In summary, starting an interleaved MRS-fMRI acquisition with a cold scanner leads to intense field drifts during the MRS acquisition that can strongly affect the GABA measurement reducing the GABA SNR, however this effect decreases after multiple acquisition sessions, as the scanner gets warmer. The field drifts have no substantial effect on the spectral quality when the acquisition is run with a non-cold scanner.

The GABA SNR trend over the first 60 averages is shown in Figure S5 F. Over this window we have the steepest SNR increase[10], and negligible field drift with non-cold scanner.

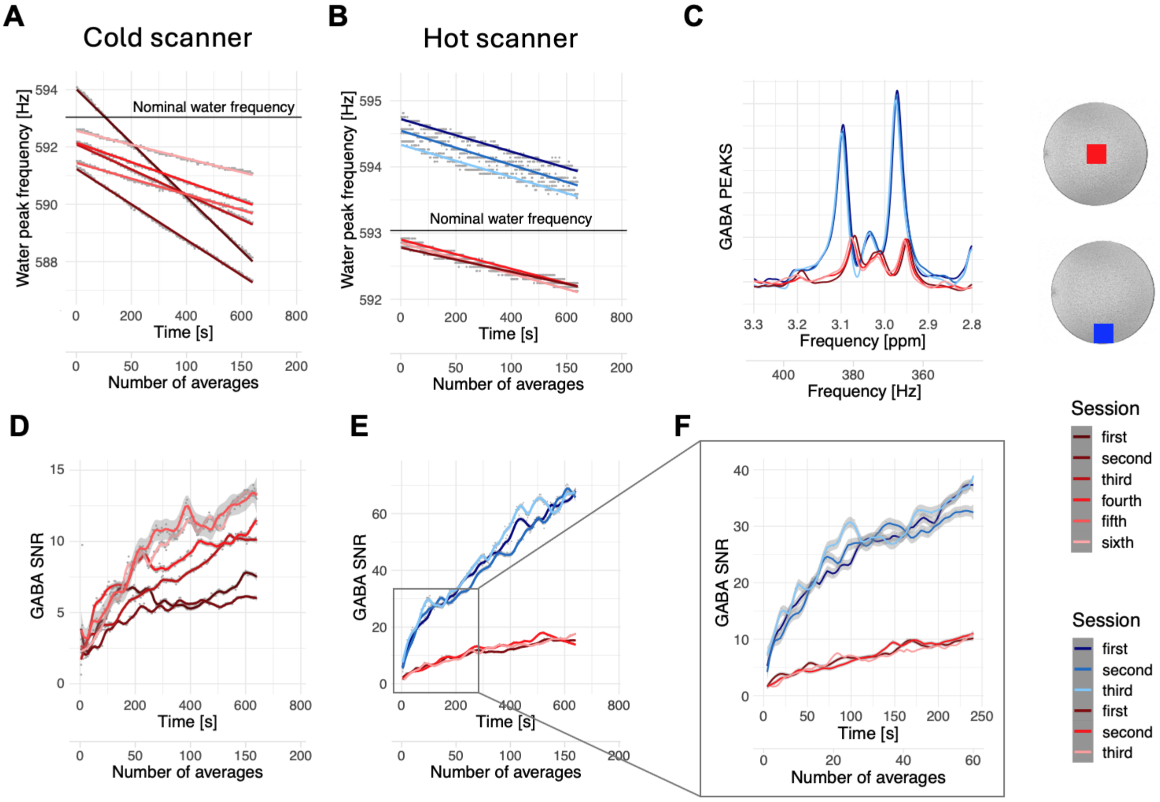


Figure S5: Effect of interleaving fMRI and MRS (phantom experiment). Panels A, C and D respectively show the water peak drift, the GABA peaks and the GABA signal to noise ratio (SNR) for a 6-sessions interleaved experiment (started with the fMRI and cold scanner). Panels B,C and E show the same results for a 3-sessions interleaved experiment (started with a non-cold scanner). In this case the voxel was positioned in the center of the spherical phantom (in red) and near the phantom's surface (in blue). Panel F shows a zoom on the first 60 averages of the GABA SNR plot in panel E. The nominal water frequency is set to 593 Hz.

##### 1.2.4 Edited fMRS quality assurance

Following data exclusion criteria (see Main Text, Material and Methods, MRS Quality control) we assessed the improvement in MRS data quality in the experimental sample (N=12) compared to the full sample (N=24). To measure the gain in MRS spectral quality we considered the following: (i) mean water frequency shift from the nominal value, (ii) within-session water frequency drift and (iii) overall edited spectral quality. Criteria (i) and (ii) are reported in Figure S6. Panel a shows a sensible reduction in average water frequency shift in the four sessions (cognitive loadings), panel b shows the within-session frequency drift of water peak as a function of time. Criterion (iii) is depicted in Figure S7: it shows reduced variability in edited spectra within-condition in the experimental sample. Improvements in Glx and GABA peaks fitting is visible in all four conditions, as confirmed by quantitative evaluation of SNR, Fit Error and FWHM (Table S2).

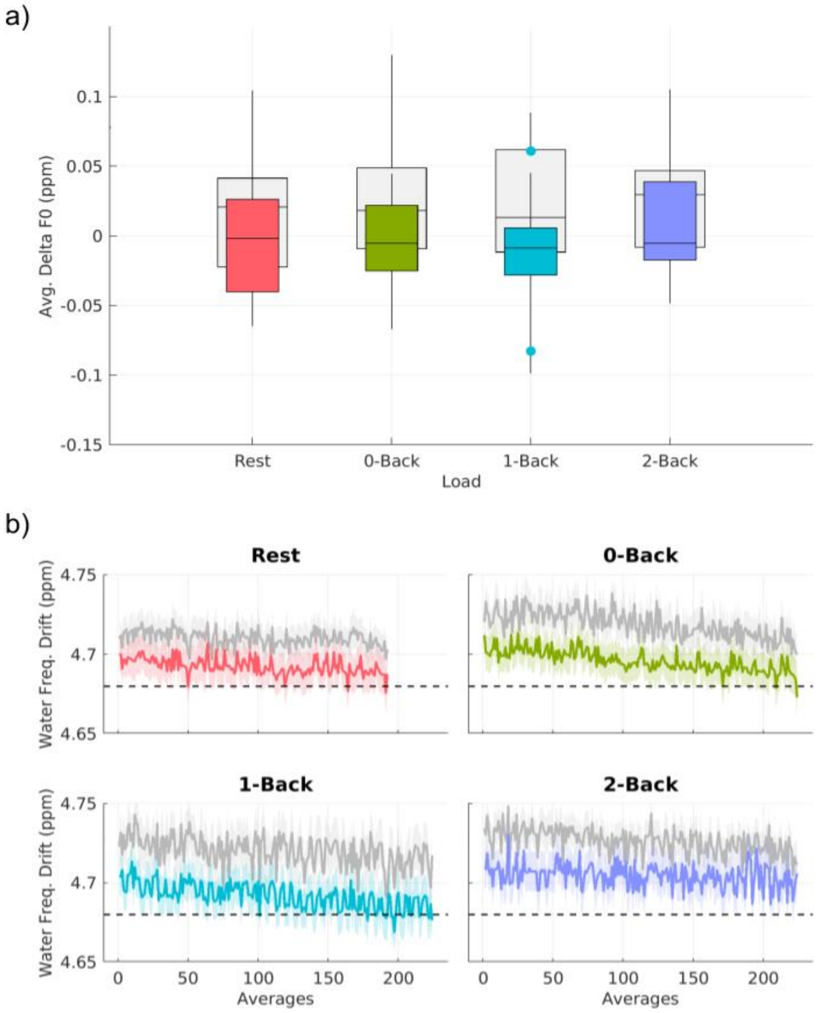


Figure S6: **Water reference drift for detecting goodness of shimming.** Panel a) FWHM of water peak determined on the entire population (grey boxes) and in the experimental subset considered in the study (colour boxes) in the four loading conditions. Panel b) Time-dependent drift of water reference peak in the entire population (grey lines) and in the experimental subset considered (colour lines). Each colour represents the four different cognitive loadings. In both cases, QA criteria were beneficial in improving the quality of fMRS data considered in the study.

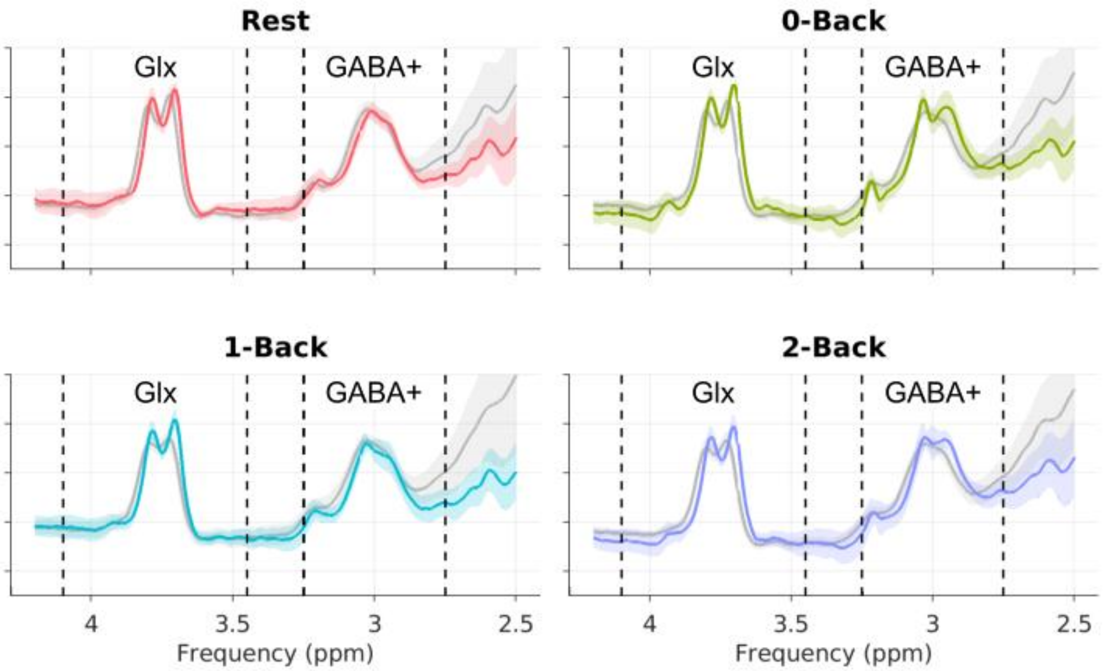


Figure S7: **Spectral quality of GABA-edited MRS fitted in Gannet.** The four panels show the improvement of Glx and GABA+ fitted spectra on the entire population (N=24, grey line) and on the experimental subset after QA (N=12, colour line) for the different cognitive loadings. Dashed vertical lines indicate frequency bands for Glx and GABA+ (3.45-4.1 and 2.75-3.25ppm, respectively).

| **QA Metric** *(mean ± SD)* | **Sample Size** *N = 24* | *$N_{QA}$=12* |
|---|---|---|
| GABA SNR | 33.2±46.3 | 22.1±3.5 |
| GABA Fit error | 9.1±10.3 | 7.0±3.0 |
| GABA FWHM | 21.8±28.4 | 19.4±2.3 |
| Glx SNR | 26.6±7.8 | 26.4±3.9 |
| Glx Fit error | 8.8±22.1 | 5.9±2.5 |
| Glx FWHM | 16.8±6.0 | 15.6±3.0 |

Table S2: **Data quality of GABA-edited MRS fitted in Gannet.** The first column is showing data quality in the whole sample (N=24), and data quality improvement after removal of outliers ($N_{QA}$=12).

#### 1.3 Edited fMRS Preprocessing

Preprocessing of in-vivo edited fMRS spectra was implemented using all the standard tools available in Gannet Gannet v3.2.0 (https://github.com/markmikkelsen/Gannet.git), and include: (i) eddy current correction both for water and metabolite data; (ii) phase correction; (iii) zero-filling of FIDs; (iv) FIDs apodization; (v) removal of residual water signal from edited spectra; (vi) spectral alignment; (vii) weighted averaging of transients.

### Supplementary Results

#### 2.1 Behavioral performance tests

The Shapiro-Wilk test showed that the distribution of accuracy (fMRI: 1-back, W = 0.70, p-value < 0.01; 2-back, W = 0.88, p-value < 0.01; fMRS: 1-back, W = 0.62, p-value < 0.001; 2-back, W = 0.90, p-value = 0.2), *d'* (fMRI: 0-back, W = 0.57, p-value < 0.001; 1-back, W = 0.82, p-value < 0.001; 2-back, W = 0.97, p-value = 0.5; fMRS: 0-back, W = 0.61, p-value < 0.001; 1-back, W = 0.85, p-value = 0.04; 2-back, W = 0.93, p-value = 0.4), and RT (fMRI: 0-back, W = 0.79, p-value < 0.001; 1-back, W = 0.86, p-value < 0.001; 2-back, W = 0.92, p-value < 0.001; fMRS: 0-back, W = 0.84, p-value < 0.001; 1-back, W = 0.84, p-value < 0.001; 2-back, W = 0.92, p-value < 0.001) values significantly departed from normality in most cases.

Despite performance accuracy reaching ceiling effect in both fMRI (0-back: 100%, 1-back: 61%, 2-back: 19%) and fMRS (0-back: 100%, 1-back: 73%, 2-back: 30%) blocks, the planned Friedman test revealed that, with increased WM load, accuracy (fMRI: $\chi 2(2) = 37.62$, p-value < 0.001; fMRS: $\chi 2(2) = 12.1$, p-value < 0.001, see Table 2) and *d'* (fMRI: $\chi 2(2) = 44.83$, p-value < 0.001; fMRS: $\chi 2(2) = 12.05$, p-value < 0.001, see Table 2) significantly decreased. Moreover, the generalized linear model showed that the WM task had a significant effect on RTs (fMRI: Run1-back, estimate $\beta = 0.12$, p-value < 0.001; Run2-back, estimate $\beta = 0.37$, p-value < 0.001; fMRS: Run1-back, estimate $\beta = 0.05$, p-value = 0.04; Run2-back, estimate $\beta = 0.29$, p-value < 0.001, see Table 2), with RTs increasing as the task difficulty parametrically increased.

**<u>Table S3:</u> Behavioural performance across different sessions.**

| Behavioural metrics | Acquisition type | Working memory tasks (mean ± sd) | | | Post hoc pairwise comparisons | |
|---|---|---|---|---|---|---|
| | | 0-back | 1-back | 2-back | Comparisons | p-value |
| Accuracy (%) | fMRI | 99.0 ± 4.7 | 96.4 ± 12.6 | 88.1 ± 13.5 | 0-back vs 1-back<br>1-back vs 2-back<br>0-back vs 2-back | **$3.4 \times 10^{-4}$**<br>**$1.7 \times 10^{-5}$**<br>**$1.1 \times 10^{-9}$** |
| | fMRS | 100 ± 0.0 | 98.2 ± 3.4 | 94.9 ± 4.9 | 0-back vs 1-back<br>1-back vs 2-back<br>0-back vs 2-back | 0.1970<br>0.1364<br>**0.0027** |
| d’ | fMRI | 4.9 ± 0.4 | 4.4 ± 0.9 | 3.6 ± 0.8 | 0-back vs 1-back<br>1-back vs 2-back<br>0-back vs 2-back | **0.0012**<br>**$1.4 \times 10^{-5}$**<br>**$9.0 \times 10^{-11}$** |
| | fMRS | 5.0 ± 0.2 | 4.7 ± 0.6 | 4.0 ± 0.8 | 0-back vs 1-back<br>1-back vs 2-back<br>0-back vs 2-back | 0.2420<br>0.1108<br>**0.0038** |
| RT (ms) | fMRI | 518 ± 123 | 586 ± 169 | 753 ± 264 | 0-back vs 1-back<br>1-back vs 2-back<br>0-back vs 2-back | **< 0.0001**<br>**< 0.0001**<br>**< 0.0001** |
| | fMRS | 544 ± 124 | 584 ± 152 | 759 ± 249 | 0-back vs 1-back<br>1-back vs 2-back<br>0-back vs 2-back | 0.0884<br>**< 0.0001**<br>**< 0.0001** |

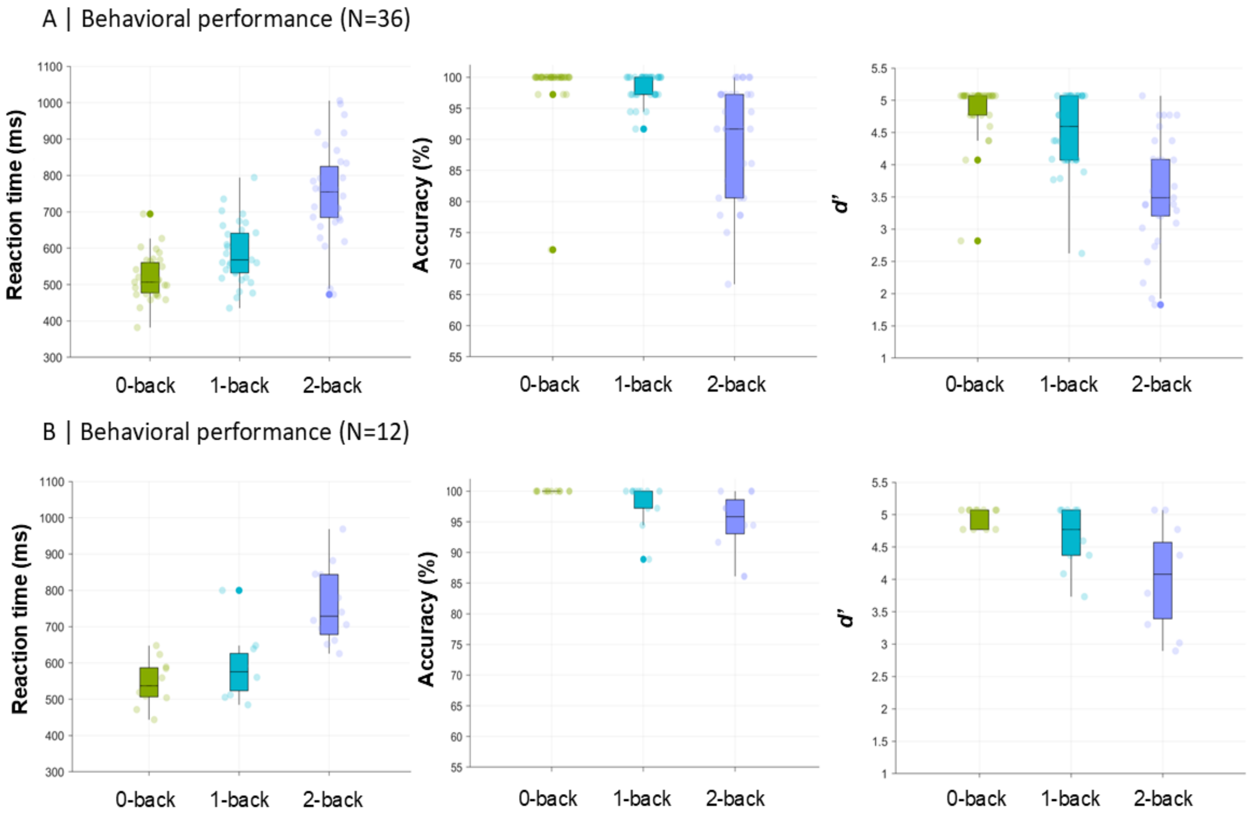


Figure S8: Boxplots of behavioural performance. Panel A) shows differences in Reaction times (RT, ms) across different runs. Panel B) shows differences in total accuracy (%) across different runs. Panel C) shows differences in total accuracy for letters stimuli (%) across different runs. Panel D) shows differences in total accuracy for numbers stimuli (%) across different runs. *=p-value FDR<0.05

##### 2.1.1 The effect of executive networking on behavioural performance

Investigating the interaction between FPN temporalities and behavioural performance for the MRS sample, we found a significant effect only for accuracy; particularly for resilience interaction with session (2-Back) ($\beta = -1.48$, $p < 0.05$, FDR corrected; Figure S9, Panel B). The lack of additional significant effects is likely due to the constrained statistical power associated with the small sample size.

##### 2.1.2 The effect of EIB on behavioural performance

Investigating the interaction between FPN temporalities and behavioral performance for the MRS sample (whole-sample described in main manuscript), we found a significant effect all three behavioral measures. Particularly, we found effects for in-degree interaction with session (2-Back) (accuracy: $\beta = -0.11$, $p < 0.05$, FDR corrected; RT: $\beta = 0.36$, p-value$_{FDR} < 0.05$; *d'*: $\beta = -0.32$, p-value$_{FDR} < 0.05$), out-degree interaction with session (2-Back) (accuracy: $\beta = -0.10$, p-value$_{FDR}$ < 0.05; RT: β = 0.37, p-value$_{FDR}$ < 0.05; *d'*: β = -0.36, p-value$_{FDR}$ < 0.05), resilience interaction with session (2-Back) (accuracy: β = -0.10, p-value$_{FDR}$ < 0.05; RT: β = 0.28, p-value$_{FDR}$ < 0.05; *d'*: β = -0.23, p-value$_{FDR}$ < 0.05), betweenness centrality interaction with session (2-Back) (accuracy: β = -0.10, p-value$_{FDR}$ < 0.05; RT: β = 0.35 p < 0.05, FDR corrected; *d'*: β = -0.24, p-value$_{FDR}$ < 0.05) and occurrences interaction with session (2-Back) (accuracy: β = -0.12, p-value$_{FDR}$ < 0.05; RT: β = 0.31, p-value$_{FDR}$ < 0.05; *d'*: β = -0.33, p-value$_{FDR}$ < 0.05).

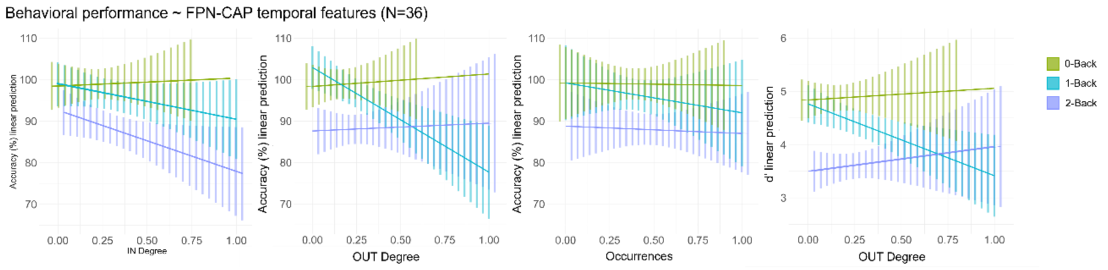


Figure S9: Relationship between behavioural performance and temporal dynamics of executive functional networks. Panel A) Illustrates the relationship between accuracy (%) and the resilience property of the FPN CAP, highlighting interactions across runs in the fMRS subgroup. Panel B) shows the association between behavioural performance (accuracy (%) and *d'*) and the temporal properties of FPN CAP in the whole sample.

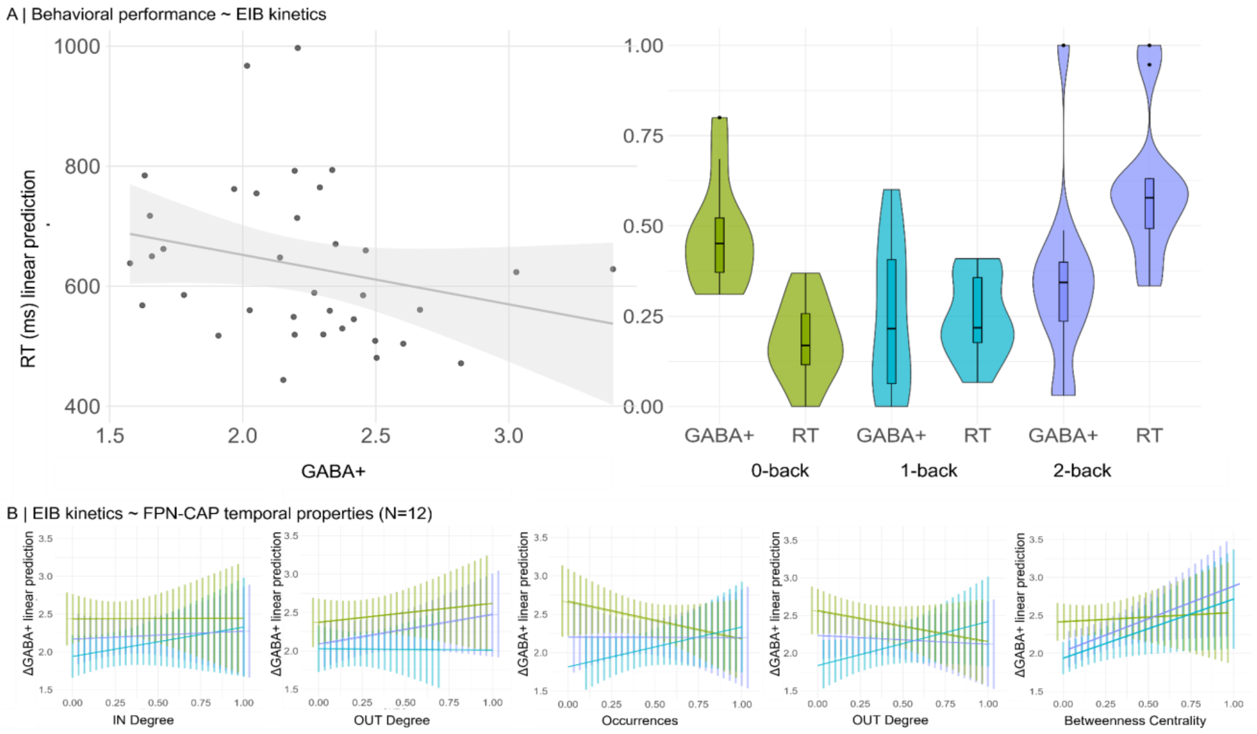


Figure S10: Relationship between behavioural performance, temporal dynamics of executive functional networks and EIB kinetics. Panel A) shows the fixed effects of static GABA+ concentration on Reaction Times (RT), while the accompanying boxplot presents the distribution of these variables across runs. Panel B) displays the association between static GABA+ concentration, expressed as delta GABA+ (the resting state value subtracted as baseline), and the temporal properties of FPN CAP.

#### 2.2 Co-activation patterns analysis

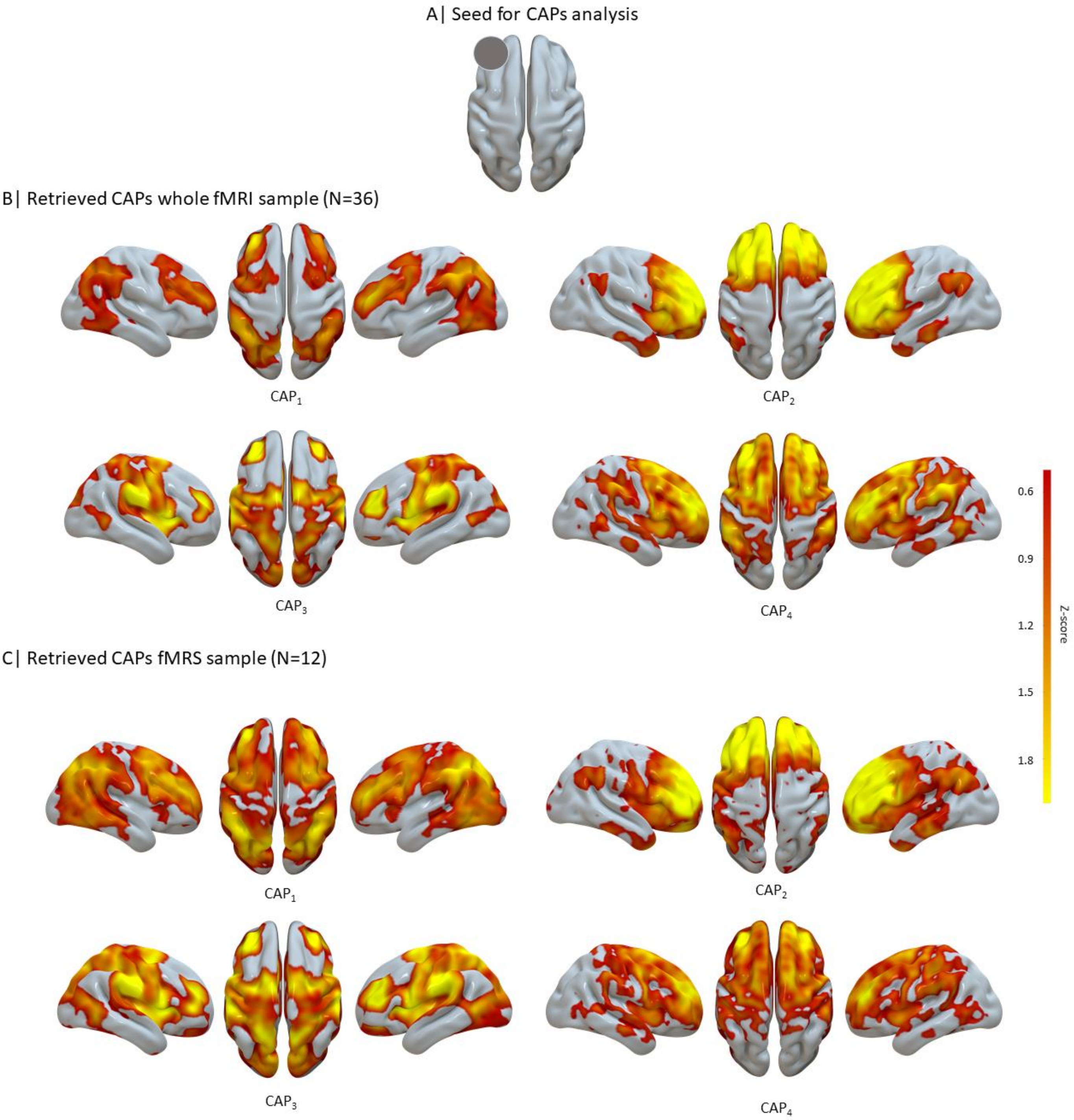


Figure S11: **Retrieved co-activations pattern (CAPs)**. A) CAPs Seed location in left DLFPC. B) Retrieved CAPs in the whole sample (N=36) by concatenating different runs. C) Retrieved CAPs in the fMRS sample (N=12) by concatenating different runs.

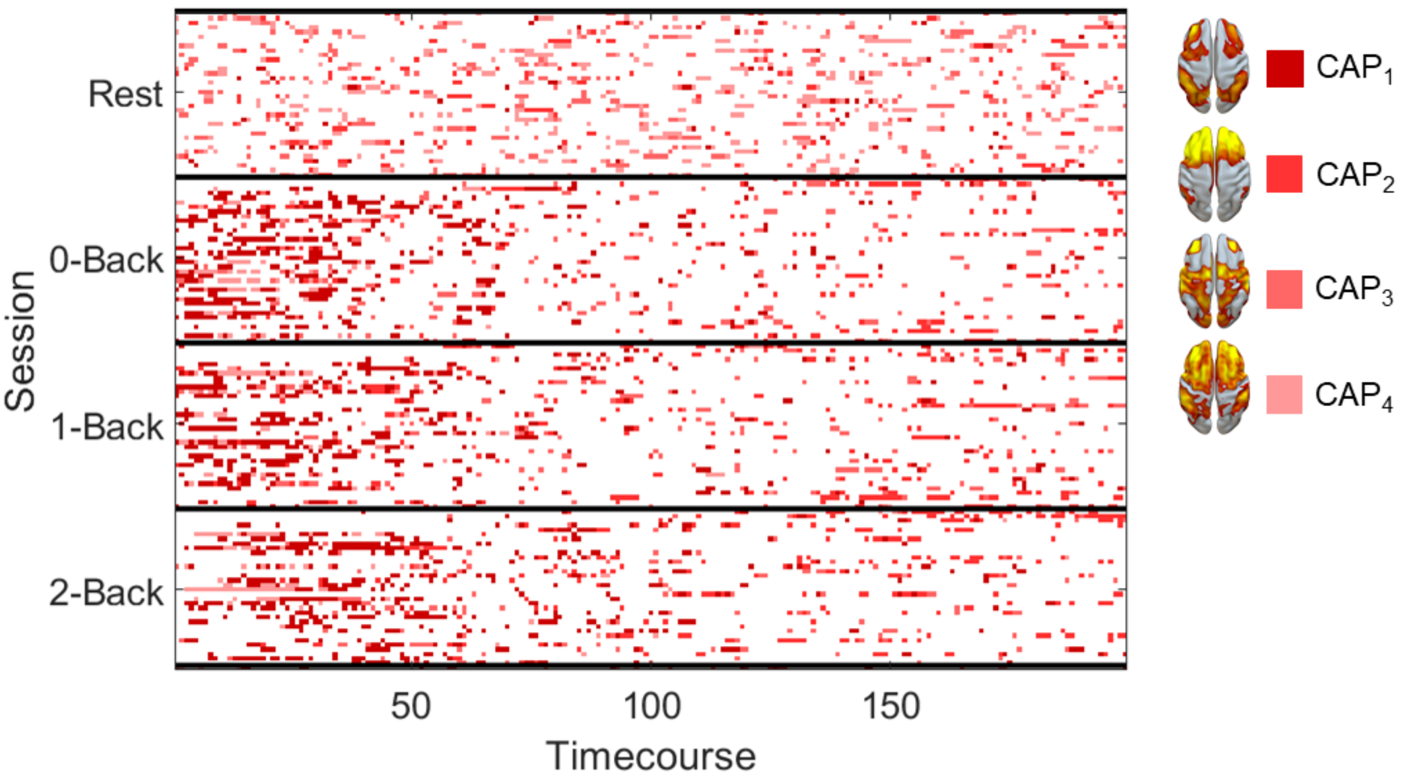


Figure S12: Co-activations pattern traces in different runs for the whole sample (N=36).

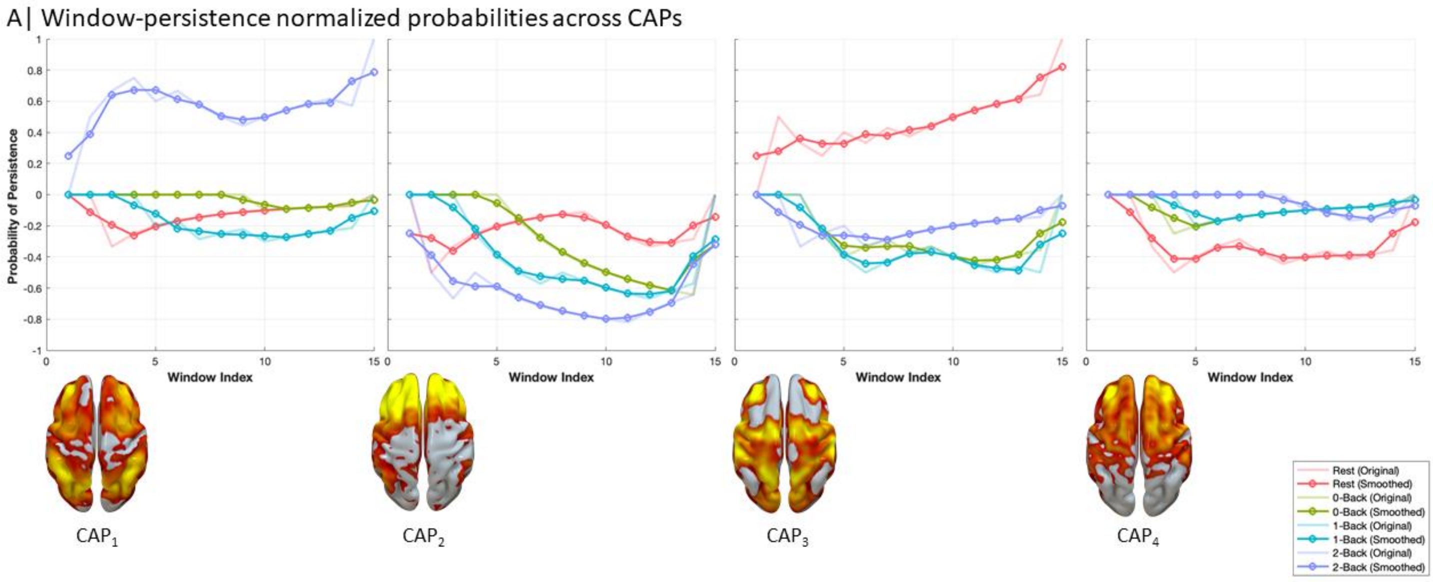


Figure S13: Normalized window-persistence probabilities of Co-activations patterns (CAPs) in different runs for the MRS sample (N=12).

**Table S4:** **Temporal properties of FPN different sessions in MRS sample.**

| Temporal FPN properties | Working memory conditions | | | | Differences across conditions (FDR corrected) | | |
|---|---|---|---|---|---|---|---|
| | rest | 0-back | 1-back | 2-back | Post-hoc comparisons | p-value | Chi-square/ A-B estimate |
| **Occurrences (%)** | 8.4±7.6 | 37.9±21.7 | 38.9±24.3 | 45.5±25.7 | | **<0.001**** | **18.24** |
| | | | | | **0-back>rest** | **<0.01*** | **18.58** |
| | | | | | **1-back>rest** | **<0.01**** | **18.95** |
| | | | | | **2-back>rest** | **<0.001**** | **21.62** |
| **Resilience** | 0.002±0.003 | 0.03±0.03 | 0.03±0.02 | 0.04±0.03 | | **<0.01*** | **15.67** |
| | | | | | **0-back>rest** | **<0.05*** | **16.75** |
| | | | | | **1-back>rest** | **<0.01*** | **18.75** |
| | | | | | **2-back>rest** | **<0.01**** | **18.66** |
| **Betweenness centrality** | 0.1±0.3 | 0.5±0.8 | 0.3±0.8 | 0.6±0.8 | | >0.05 | |
| **IN Degree** | 0.004±0.004 | 0.008±0.007 | 0.006±0.008 | 0.007±0.006 | | >0.05 | |
| **OUT Degree** | 0.003±0.004 | 0.007±0.008 | 0.006±0.006 | 0.007±0.007 | | >0.05 | |

#### 2.3 Static edited fMRS

Time integral of the EIB ratio was computed by averaging the entire set of edited spectra to investigate the global effect of the cognitive task on metabolites of interest. Figure SX, panel a) illustrates the absence of relevant changes in EIB as a function of WM load; these findings are supported by non-significant differences across runs. Despite this, a qualitative trend towards an increase of Glx, rather than GABA, as reported in Fig Sx, panel b), is consistent with results obtained from time-varying analysis.

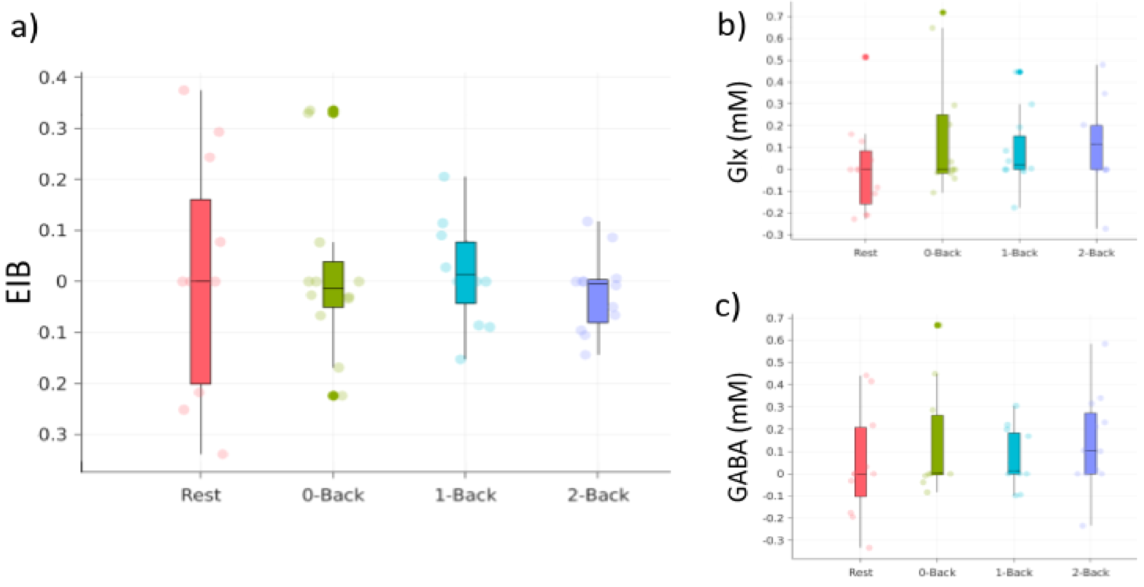


Figure S14: **Boxplots of metabolites concentrations across runs.** EIB ratio, Glx and GABA+ concentrations with alpha tissue correction (panels a, b, and c, respectively) were not significantly different across runs using one-way ANOVA.

#### 2.4 Dynamic edited fMRS

##### 2.4.1 Temporal properties

Temporal dynamics of EIB and metabolites were evaluated by the following metrics: (i) Area under the curve (AUC), (ii) Out Degree and (iii) Kullback-Leiber divergence with a Kruskal-Wallis nonparametric test. Results for (i) are reported in the main text.

A significant effect of *Load* (Chi-squared(3,44)=9.92, p-value=0.02, Figure S14) for EIB Out Degree was found. Post-hoc comparisons showed that the 1-back condition had a significantly higher EIB Out Degree relative to resting-state condition (rest < 1-Back, p-value=0.02, Mean Ranks Difference=-16.5, Figure S14). Similarly, a significant effect of Out Degree for GABA was reported (Chi-squared(3,44)=8.24, p-value=0.04, Figure S14).On the other hand, for what concerns EIB, Kullback-Leiber divergence showed no significant effects. Whereas for Glx and GABA a significant effect of *Load* was found (Glx: Chi-squared(3,44)=9.71, p-value=0.02, Figure S14; GABA: Chi-squared(3,44)=10.44, p-value=0.01). Post-hoc comparison revealed an increase for Glx stationarity in 1-back (rest<1-Back, p-value=0.02, Mean Ranks Difference=-16.0) and for GABA stationarity in 0-back (rest<0-Back, p-value=0.02, Mean Ranks Difference=-16.1).

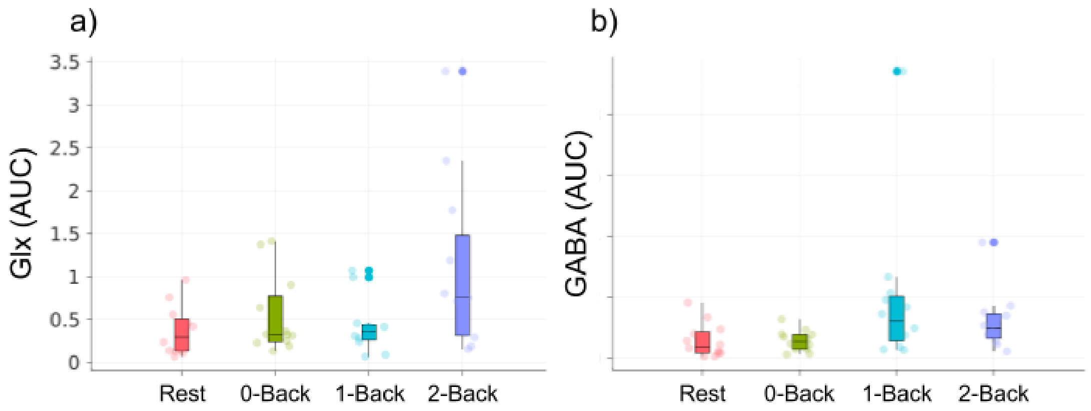


Figure S15: **Area under the curve (AUC) for Glx, panel a) and GABA+, panel b).** A significant effect of GABA+ is reported (main effect, p-value=0.01); no differences were found for Glx. See main text for relevant details.

<u>Table S5:</u> Time-series conventional and graph-properties of EIB dynamics.

| **Time-series properties** | **Cognitive load session** ($N_{QA}$=12) | | | |
|---|---|---|---|---|
| | Rest | 0-back | 1-back | 2-back |
| Slope (mean±SD) | 0.001±0.1 | 0.045±0.1 | 0.006±0.2 | 0.109±0.2 |
| Time-to-peak (frames mean±SD) | 10.1±3.2 | 14.2±5.3 | 13.5±4.5 | 12.5±5.3 |
| Zero-crossing (mean±SD) | 1.8±0.7 | 2.3±1.5 | 2.2±1.7 | 1.8±1.5 |
| AUC (mean±SD) | 0.49±0.40 | 0.64±0.42 | 1.63±1.55 | 1.88±1.82 |
| Out-degree (mean±SD) | 1.1±0.2 | 1.3±0.2 | 1.4±0.2 | 1.3±0.2 |
| Kullback-Leiber divergence (mean±SD) | 0.003±0.006 | 0.02±0.03 | 0.03±0.03 | 0.04±0.06 |

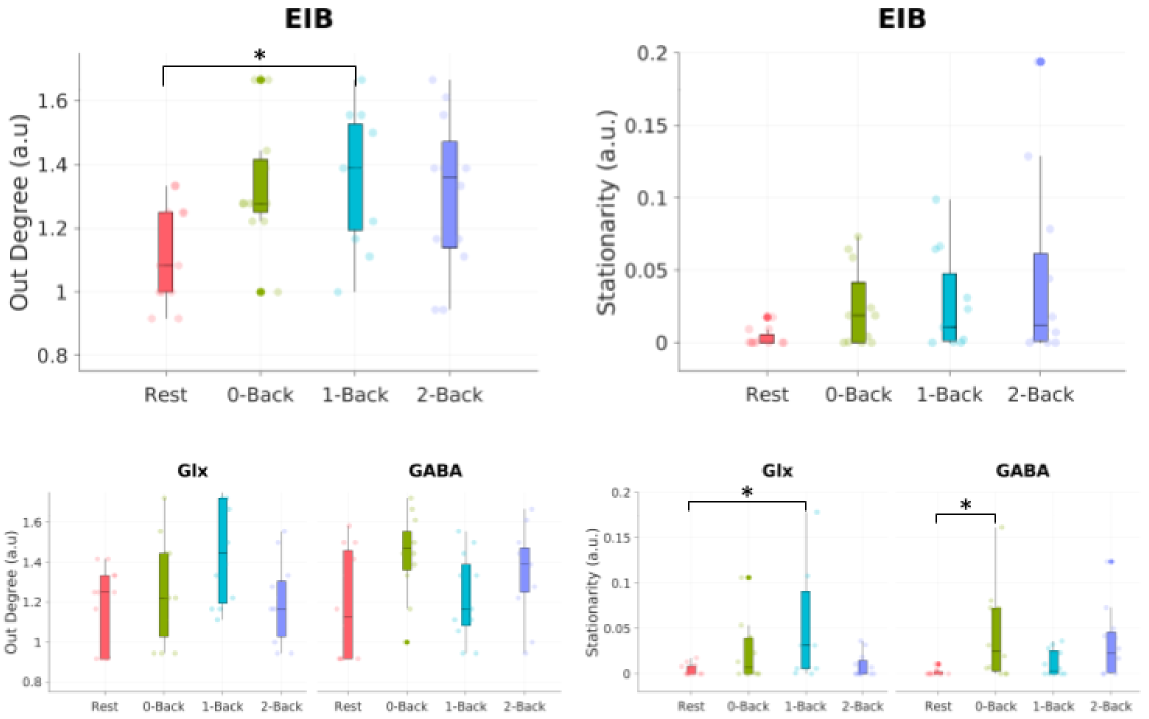


Figure S16. **Temporal properties of EIB ratio.** *Upper row*: Out degree and Kullback-Leiber divergence (KLD) extracted from EIB dynamic analysis. The two metrics are graph-theoretical proxies for slope and stationarity. A significant increase of out degree is reported between Rest and 1-Back ($p=0.02$). Concerning stationarity, only a positive trend is reported with no statistical evidence. However, increasing mean values of KLD reflect a time variant behavior of the EIB curve, with lower stationarity over time depending on the WM load. *Bottom row*: Out degree and KLD of separate Glx and GABA components. Asterisks mark significant differences, $p<0.05$.
